# The laminin-keratin link shields the nucleus from mechanical deformation and signalling

**DOI:** 10.1101/2022.03.01.482474

**Authors:** Zanetta Kechagia, Pablo Sáez, Manuel Gómez-González, Martín Zamarbide, Ion Andreu, Thijs Koorman, Amy E.M. Beedle, Patrick W.B. Derksen, Xavier Trepat, Marino Arroyo, Pere Roca-Cusachs

## Abstract

The mechanical properties of the extracellular matrix (ECM) dictate tissue behaviour. In epithelial tissues, laminin is both a very abundant ECM component, and a key supporting element. Here we show that laminin hinders the mechanoresponses of breast epithelial cells by shielding the nucleus from mechanical deformation. Coating substrates with laminin-111, unlike fibronectin or collagen I, impairs cell response to substrate rigidity, and YAP nuclear localization. Blocking the laminin-specific integrin β4 increases nuclear YAP ratios in a rigidity dependent manner, without affecting cell forces or focal adhesions. By combining mechanical perturbations and mathematical modelling, we show that β4 integrins establish a mechanical linkage between the substrate and the keratin cytoskeleton, which stiffens the network and shields the nucleus from actomyosin-mediated mechanical deformation. In turn, this affects nuclear YAP mechanoresponses and chromatin methylation. Our results demonstrate a mechanism by which tissues can regulate their sensitivity to mechanical signals.

## Introduction

Cells are subjected to continuous reciprocal interactions with the surrounding extracellular matrix (ECM). ECM parameters such as its composition or mechanical properties shape cellular responses, from development and tissue morphogenesis to tissue repair and homeostasis. Abnormal cell-ECM interactions drive pathological conditions such as cancer and fibrosis^1^, and are promising targets for therapeutic interventions. Cell membrane receptors such as integrins act as biochemical sensors that recognize specific epitopes on ECM molecules^1^. Different combinations of ECM molecules trigger intracellular events that propagate cell- and tissue-specific responses through changes in gene expression and signaling. Integrins also respond to changes in ECM mechanical properties^2^, in a process known as mechanotransduction. Increased ECM stiffness can lead to the activation, clustering, and maturation of integrin adhesions. In turn, this triggers cytoskeletal rearrangements and a build-up of intracellular tension that can propagate to the cell nucleus, where it regulates the nuclear localization and activity of transcriptional regulators such as YAP^2–4^.

This framework of integrin-mediated cell response to increased ECM stiffness and mechanotransduction to the nucleus, has largely been studied for ECM components such as fibronectin and collagen^3,5,6^, as well as for reconstituted basement membranes such as Matrigel^7–11^. However, the specific role of laminin, a common substrate for all epithelial tissues, is unclear. Laminin forms an important part of the basement membrane (BM) underlying epithelial tissues and guides pivotal cellular processes ranging from healthy epithelial homeostasis to cancer metastasis^12–18^. Changes in BM composition or mechanical properties are critical in several stages of cancer progression, regulating both tissue organization as well as tumour invasiveness^19,20^. This role of the BM has been particularly well characterized and reported for the case of breast cancer^7,21–25^. We thus sought to investigate how cells respond to increased tissue rigidity on a laminin based extracellular environment, and how these changes can influence mechanotransduction in mammary epithelial cell models. We find that laminin-111 hinders cellular mechanoresponses through its interaction with integrin β4 and the keratin cytoskeleton. By using molecular and mechanical perturbations in combination with mathematical modelling, we show that α6β4 integrins and their incorporation to hemidesmosomes mediate the mechanical engagement between a laminin ECM and the keratin cytoskeleton. In turn, this engagement stiffens the intermediate filament cytoskeleton and shields the nucleus from mechanically induced deformation, preventing nuclear mechanoresponses. Through this work, we identify a novel tool that epithelial cells can employ to tune their response to rigidity, thereby controlling mechanosensing, nuclear integrity, and chromatin methylation.

## Results

### Laminin-111 hinders cell response to rigidity

To study the role of laminin in cell mechanoresponses, we focused on the well-known breast epithelial model of MCF10A cells, and one of the main types of laminin present in breast epithelia *in vivo* and *in vitro* models, laminin-111^22,25–27^. Mechanoresponses such as the nuclear localization of YAP are abrogated by cell-cell contact and cadherin ligation, and high nuclear YAP levels are associated with E-cadherin deficient breast tumours^28–30^. To isolate the role of cell-ECM interactions from those mediated by cell-cell contact, we studied single cells. We first compared the mechanoresponses of MCF10A cells on laminin-111 (from now on referred as laminin) to those on Collagen I and fibronectin. To this end, we used polyacrylamide (PAA) gels of rigidities between 0.5 to 30 kPa, thereby encompassing the range of soft –healthy– and stiff –malignant– breast tissue^31,32^. We coated gels with laminin, collagen I or fibronectin, and first quantified the tractions that the cells exert on each condition through traction force microscopy^33,34^. We found that cells exerted much lower tractions when seeded on laminin than when seeded on collagen I or fibronectin substrates (Fig. 1a,b).

**Fig. 1.**
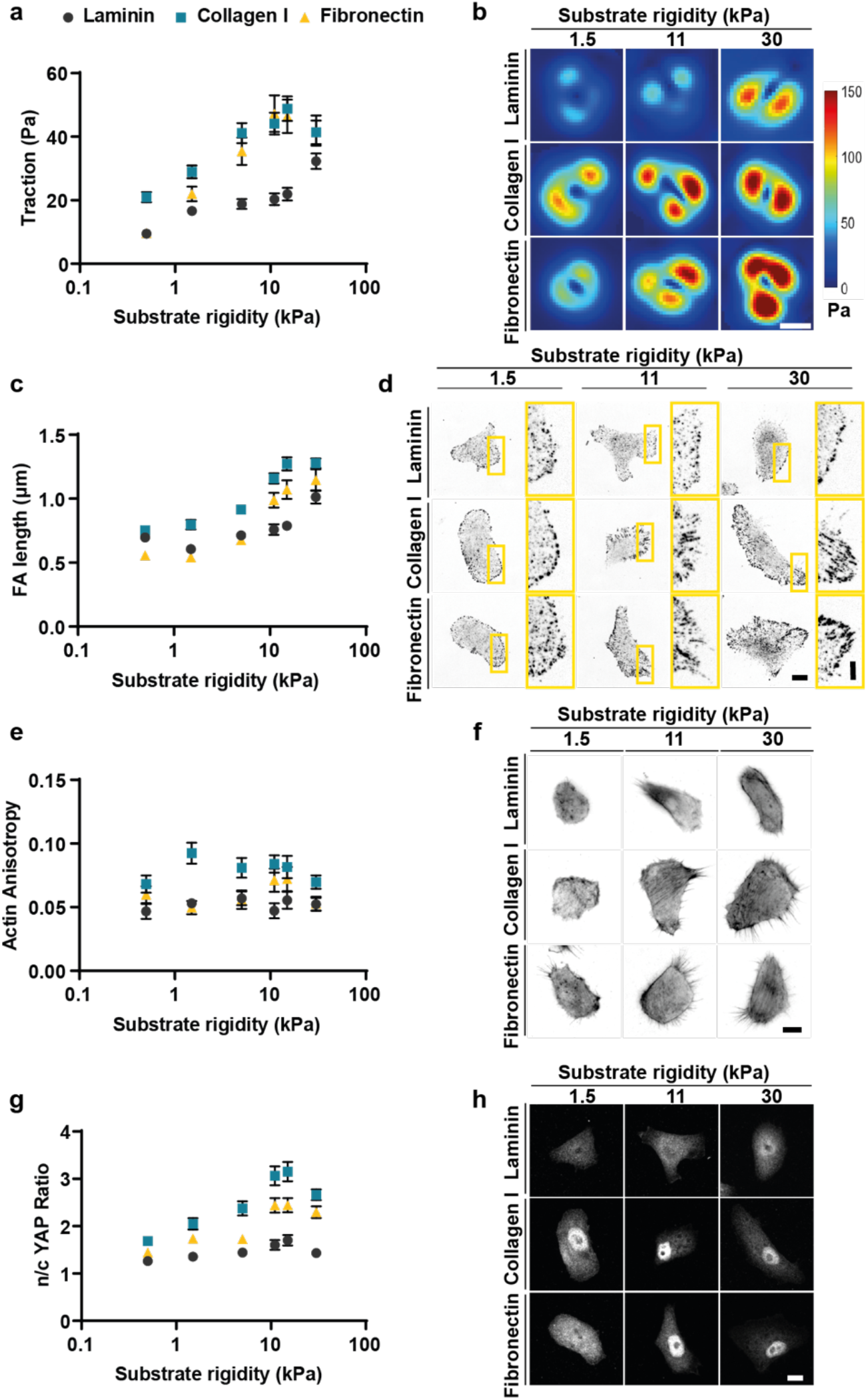
Laminin coating impairs cell mechanosensing in response to substrate rigidity. **a.** Average values of MCF10A cell tractions (*n* = 56/66/41, 59/93/45, 80/66/26, 55/50/30, 51/45/32, 79/64/23 cells for laminin/collagen/fibronectin substrates and increasing rigidity; mean of at least 3 independent experiments). The effect of both rigidity and substrate coating is significant. (*P* < 0.0001, two-way analysis of variance (ANOVA)). **b.** Corresponding example colour maps of traction forces; scale bar is 10 μm. **c.** Average values of focal adhesion length from phospho-Paxillin stainings (*n* = 69/91/48, 47/75/35, 63/89/37, 43/80/31, 53/64/52, 58/110/48 cells for laminin/collagen/fibronectin substrates and increasing rigidity; mean of at least 3 independent experiments). The effect of both rigidity and substrate coating is significant (*P* < 0.0001, two-way ANOVA). **d.** Corresponding example images of phospho-Paxillin stainings; scale bar is 10 μm. The right-hand image of each pair corresponds to the yellow rectangle in the left-hand image; scale bar is 4 μm. **e.** Quantification of actin fibre alignment (n=43/53/51, 51/52/43, 47/62/36, 54/54/30, 44/48/33, 76/50/29 cells for laminin/collagen/fibronectin substrates and increasing rigidity; mean of at least 3 independent experiments). The effect of substrate coating is significant. (*P* < 0.0001, two-way ANOVA). **f.** Corresponding images of phalloidin stainings of MCF10A cells; scale bar is 10 μm. **g.** Quantification of nuclear/cytosolic YAP ratio (*n* = 108/95/119, 60/68/80, 68/61/70, 78/75/57, 59/62/72, 123/111/84 cells for laminin/collagen/fibronectin substrates and increasing rigidity; mean of at least 3 independent experiments). The effect of both rigidity and substrate coating is significant. (*P* < 0.0001, two-way ANOVA). **h.** Corresponding example images of YAP stainings; scale bar is 10 μm. Error bars represent mean ± s.e.m.

Increased rigidity induces intracellular mechanoresponses; effects commonly observed are the growth of focal adhesions, the formation of actin stress fibres and the nuclear translocation of mechanosensitive proteins such as YAP^2^. In agreement with the lower traction force profile, cells seeded on laminin-coated PAA gels exhibited short focal adhesion length –as quantified through phosphorylated Paxillin (p-Pax) stainings–, low actin fibre alignment –as quantified by fibre anisotropy–, and low levels of YAP nuclear localization, with average nuclear to cytoplasmic (n/c) ratios below 2 (Fig. 1c-h). This phenotype was again contrasted by the cell response on collagen I and fibronectin substrates, where all parameters were higher than on laminin, and markedly increased with rigidity (Fig. 1c-h). Further validating the applicability of these results to the breast context, we found similar responses for human breast myoepithelial cells. Those cells exhibited an increase in n/c YAP ratios and FA length with rigidity (from 0.5 to 30 kPa) on collagen I or fibronectin coated substrates (Supplementary Fig. 1). In contrast, cells seeded on laminin had similar n/c YAP ratios when seeded on 0.5 and 30 kPa gels, and a small increase in FA length. Taken together, these results indicate that laminin hinders cell responses to rigidity.

### Integrin α6β4 impedes the nuclear localization of YAP

Cells interact with the ECM by forming adhesions through molecules such as integrins^2^. Specifically, adhesion to laminin is mediated mainly by α6β4, α7β1, α6β1 and α3β1 integrin dimers^35,36^. To determine their involvement in cell-laminin interactions and subsequent mechanoresponses, we blocked the corresponding integrin subunits, namely integrin β4, α6, α3 and β1, using blocking antibodies^37,38^. In all cases, focal adhesions remained small (Fig. 2a,b) in comparison to those found on collagen I or fibronectin substrates (Fig. 1c,d). However, blocking either integrin β4 or its binding partner integrin α6 had a counterintuitive result on YAP n/c ratios: rather than reducing them (as one would expect if adhesion is blocked), it increased them in a rigidity-dependent way (Fig. 2c,d), to levels similar to those found on collagen I or fibronectin substrates (Fig. 1g). Interestingly, blocking α6β4 integrins in keratinocytes has been previously reported to increase myosin phosphorylation and cell contractility, also leading to increased YAP n/c ratios^39^. However, in our case the effect of integrin α6β4 blocking was specific to nuclear YAP levels, without affecting the length of focal adhesions, traction forces (Fig. 2e,f), or phosphorylation of myosin light chain (pMLC) levels (Fig. 2g). Confirming the prominent role of nuclear YAP mechanosensing in conditions dominated by cell-matrix rather than cell-cell contacts^29,30,40^, blocking β4 integrins had a similar effect at the edge of cell monolayers, but did not affect cells in the centre of confluent monolayers, where n/c YAP ratios remained close to 1 (Supplementary Fig. 2a-c). Further confirming the mechanosensitive role of integrin β4, we found that it exhibited a rigidity dependent increase in expression (Supplementary Fig. 2d), which was absent when cells were seeded on collagen I coated substrates, suggesting that this is a laminin-specific response (Supplementary Fig. 2d).

**Fig. 2.**
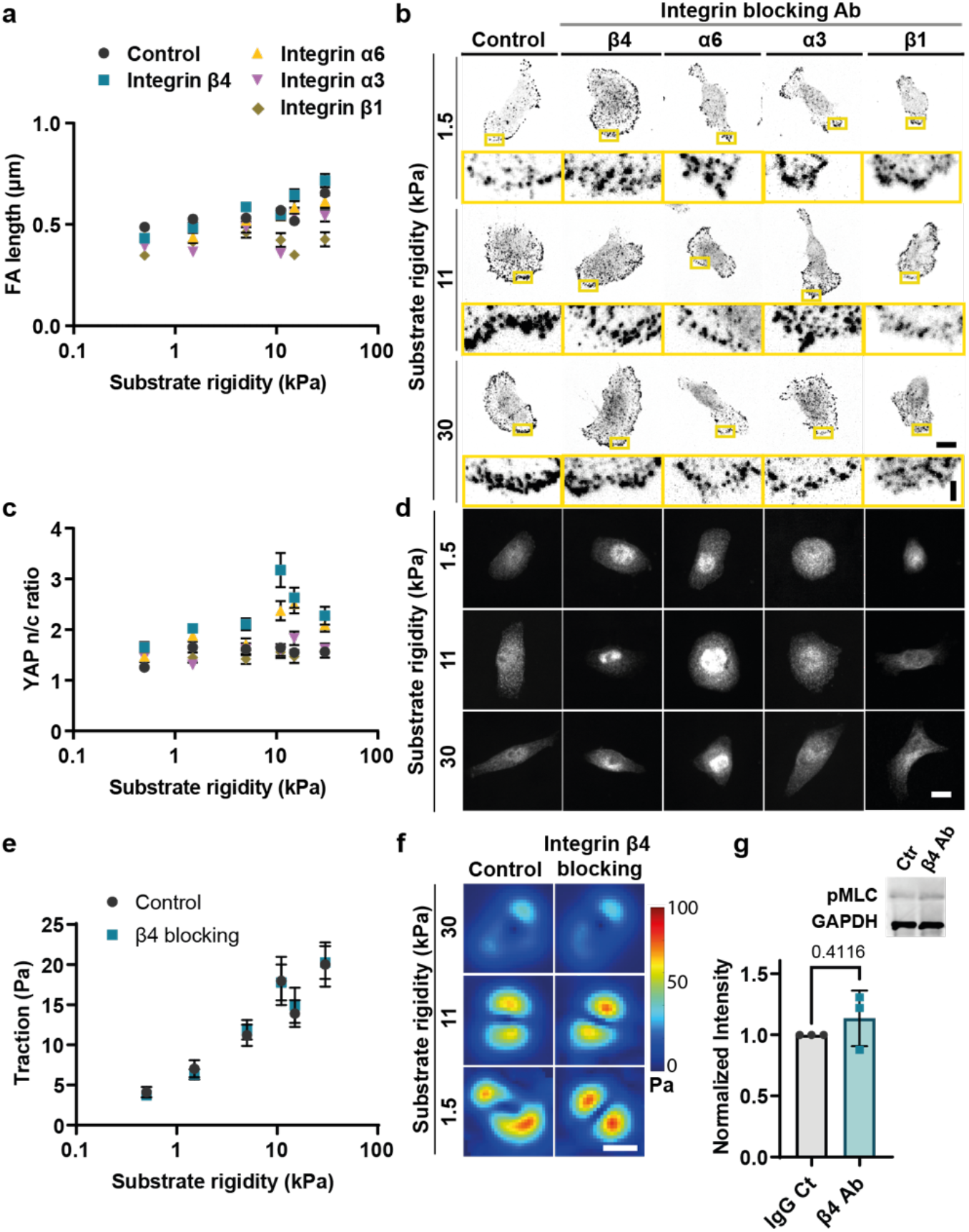
Integrin α6β4 alters nuclear mechanosensing on laminin. **a.** Average values of FA length of MCF10A cells seeded on laminin coated PAA gels of different rigidities upon treatment with different integrin blocking antibodies (*n* = 83/59/39/38/31, 94/46/46/41/37, 92/52/45/29/29, 99/53/46/24/32, 48/32/21/32/35, 62/35/21/36/29 cells treated with control/β4/α6/α3/β1 antibodies for substrates of increasing rigidity; mean of at least 3 independent experiments). The effect of both integrin blocking and substrate stiffness is significant (*P* < 0.0001, two-way ANOVA). Error bars represent mean ± s.e.m. **b.** Sample phospho-Paxillin stainings of MCF10A cells treated with control/β4/α6/α3/β1 blocking antibodies for 1.5, 11 and 30 kPa substrate stiffness; scale bars are 10 μm (main images)/2 μm (zoomed images). **c.** Average values of nuclear/cytosolic YAP ratio of MCF10A cells seeded on laminin coated PAA gels of different rigidities upon treatment with different integrin blocking antibodies (*n* = 90/61/28/30/33, 104/72/38/58/26, 101/73/49/43/35, 121/70/40/69/38, 94/51/39/51/32,105/62/51/46/34 of control/β4/α6/α3/β1 antibodies treated cells for substrates of increasing rigidity; mean of at least 3 independent experiments). The effect of both integrin blocking and substrate stiffness is significant. (*P* < 0.0001, two-way ANOVA). Error bars represent mean ± s.e.m. **d.** Sample YAP stainings of MCF10A cells treated with control/β4/α6/α3/β1 blocking antibodies for 1.5, 11 and 30 kPa substrate stiffness; scale bar is 10 μm. **e.** Average values of MCF10A cell tractions treated with control or β4 blocking antibodies (n=27/29, 27/30, 39/36, 28/30, 32/32, 28/29 of control/β4 antibody treated cells for substrates of increasing rigidity; mean of 3 independent experiments). The effect of rigidity is significant (*P*<0.0001, two-way ANOVA). Error bars represent mean ± s.e.m. **f.** Corresponding colour maps of sample single cell traction forces; scale bar is 10 μm. **g.** WB and quantification for phospho-MLC2 levels in MCF10A cells treated with control or integrin β4 blocking antibody upon normalizing to the control treated cells (n=3). No significant effect was observed, *P*>0.05, two-tailed paired t-test). Error bars represent mean ± SD.

We further examined the effect of integrin β4 blocking in the human myoepithelial cell line. Similar to MCF10A cells, blocking integrin β4 and not integrin β1 or α3 resulted in higher nuclear YAP levels for cells cultured on 30kPa PAA gels (Supplementary Fig. 2e,g). Accordingly, integrin β4 blocking did not affect focal adhesion length (Supplementary Fig. 2f,g). Overall, these results demonstrate that α6β4 integrins impede a rigidity-depended increase in the nuclear localization of YAP, without affecting cellular contractility.

### Integrin α6β4 regulates YAP mechanosensing by linking laminin to the keratin cytoskeleton via its interaction with plectin

Integrin α6β4 forms an important component of hemidesmosomes, which are epithelial cell contact sites linking the intermediate filament (IF) cytoskeleton to the BM^41^. This interaction is mediated by the binding of IFs to the intracellular domain of integrin β4 through the cytolinker plectin (Fig. 3a)^41–43^. To determine the effect of integrin β4 blocking in the morphology and cohesion of IFs, we stained for keratin 8, one of the main types of IF expressed by MCF10A cells^44^. We found that upon blocking integrin β4, the distribution of keratin 8 at the cell periphery was less homogeneous (as quantified by the coefficient of variation of the signal) and had lower intensity (Fig. 3b-d), whereas the total levels of keratin 8 intensity remained similar (Supplementary Fig. 3a). Often changes in the organization of keratin networks are indicative of altered mechanical properties^45,46^. We thus hypothesized that, since β4 does not affect traction forces or actomyosin contraction, its role in YAP nuclear localization could be mediated by changes in keratin organization.

**Fig. 3.**
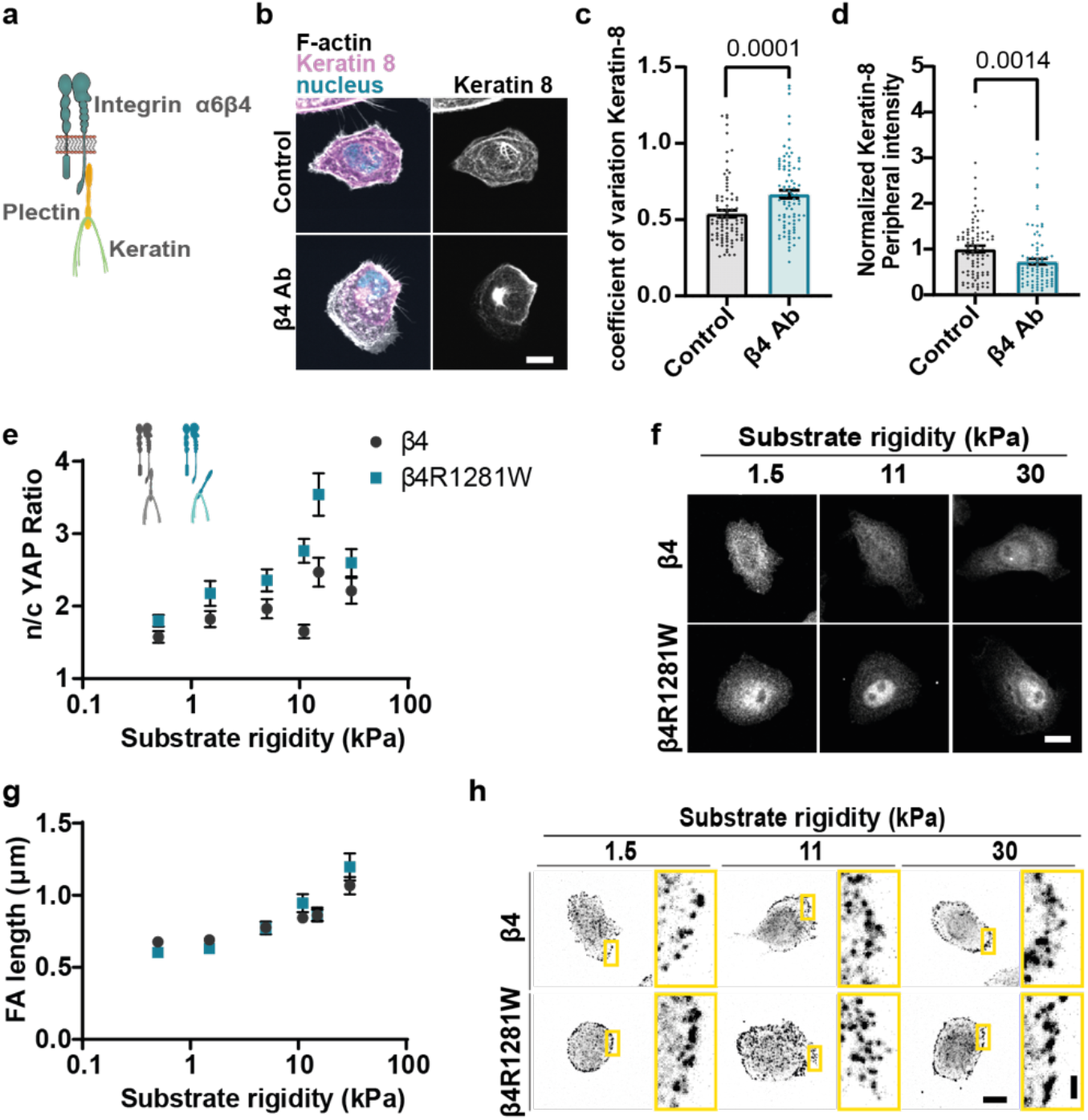
The mechanical role of integrin β4 is mediated by its connection to plectin and keratins. **a.** Schematic representation of the interaction of integrin α6β4 with plectin and intermediate filaments. **b.** Actin and Keratin 8 stainings of MCF10A cells on 11kPa PAA gels treated with IgG (control) or integrin β4 blocking (β4 Ab) antibodies; scale bar is 20 μm. **c.** Quantification of coefficient of variation of keratin 8 signal at the basal layer of the cell periphery for control or integrin β4 blocking (β4 Ab) conditions (*P*=0.0001, unpaired two-tailed t-test, n=88/87 cells for control/integrin β4 blocking conditions, for 6 independent experiments). **d.** Quantification of average mean intensity of keratin 8 signal at the basal layer of the cell periphery for control or integrin β4 blocking (β4 Ab) conditions (*P*=0.0014, two-tailed Mann-Whitney test, n= 88/87 cells for control/integrin β4 blocking conditions, for 6 independent experiments). **e.** Quantification of average n/c YAP ratios of MCF10A cells transfected with wild type β4 or mutant integrin β4R1281W (*n* = 69/65, 68/58, 63/65, 84/66, 51/52, 47/66 cells for β4 and β4R1281W and increasing rigidity; mean of 4 independent experiments). The effect of both rigidity and β4 mutation is significant. (*P*< 0.0001, two-way ANOVA). **f.** Corresponding YAP stainings of MCF10A wild type β4 or mutant integrin β4R1281W cells; scale bar is 10 μm. **g.** Quantification of FA length of MCF10A cells transfected with wild type (β4) or mutant integrin β4R1281W (*n* = 40/33, 33/35, 31/35, 39/22, 27/19, 36/36 cells for β4 and β4R1281W and increasing rigidity; mean of 3 independent experiments). The effect of rigidity is significant. (*P*< 0.0001, two-way ANOVA). **h.** Corresponding pPAX stainings of MCF10A wild type (β4) or mutant integrin β4R1281W cells; scale bars are 10 μm (main images)/2 μm (zoomed images). Error bars represent mean ± s.e.m.

Integrin β4, aside from being a structural component of hemidesmosomes, is also a major signaling hub^47^. To examine the effect of integrin β4-keratin interactions without affecting its other functions or membrane localization, we introduced an integrin β4 mutant (integrin β4R1281W) that has lower affinity for plectin, and therefore for IFs (Supplementary Fig. 3b,c)^42,43^. As a control, we transfected MCF10A cells with the wild type version of integrin β4-GFP (Supplementary Fig. 3b,c). As expected, cells expressing mutant integrin β4R1281W lacked hemidesmosome structures (Supplementary Fig. 3d). This was observed by imaging either the transfected mutated integrin, or the endogenous WT integrin, suggesting a dominant negative effect of the mutation (Supplementary Fig. 3d). We next examined the effect of integrin β4-keratin interaction on the subcellular localization of YAP. Similar to previous results obtained by integrin β4 blocking antibodies, MCF10A cells expressing the mutant integrin β4R1281W had a higher nuclear to cytoplasmic YAP ratio that was rigidity dependent, peaking around 11-15kPa (Fig. 3e,f). As before, the length of focal adhesions and tractions were similar between the two cell types (Fig. 3g,h and Supplementary Fig. 3e,f). Thus, the effect of β4 integrins in the nuclear localization of YAP is mediated by the anchorage between keratins and the laminin substrate.

### Keratin-substrate interactions alter cytoskeletal dynamics and organization

To explore the mechanism involved in laminin-mediated effects in nuclear mechanoresponses, we first hypothesized that the keratin cytoskeleton could directly regulate force transmission to the nucleus, which is known to trigger YAP nuclear localization^3^. To assess this, we knocked down nesprin-3, which connects the keratin cytoskeleton to the nuclear lamina^48^(Supplementary Fig. 3g,h). Nesprin-3 depletion strongly impacted cell phenotype, leading to cell rounding, perinuclear collapse of the actin and keratin-8 cytoskeleton, as well as low YAP n/c ratios (Supplementary Fig. 3i-k). This effect was opposite to that induced by blocking α6β4 or its connection to keratins. Thus, a direct link between keratins and the nucleus via nesprin-3 does not explain our results, although it highlights that the interaction between the keratin cytoskeleton and the nucleus is important in maintaining cellular integrity.

Alternatively, the keratin cytoskeleton could be affecting the nucleus in a less direct way, by regulating how actin-mediated force generation reaches the nucleus. Indeed, the actin and intermediate filament cytoskeletal networks are tightly interconnected^49,50^, and keratin filaments undergo retrograde flows driven by actomyosin contractility^51–53^. To explore how such a mechanism would operate, we implemented a computational model for the actomyosin and keratin cytoskeletons (see Methods). The model considers the actomyosin cytoskeleton as an active and viscous gel undergoing turnover, in which myosin contractility leads to a continuous flow of actin towards the cell centre (retrograde flow)^54,55^. The keratin cytoskeleton is modelled as a passive viscoelastic gel undergoing a slower turnover as compared to actin^56–58^. The actomyosin network drags the keratin cytoskeleton via an inter-network friction, leading also to keratin retrograde flow. Each network is in turn connected to the underlying substrate through integrin-mediated adhesions, which we model with cytoskeletal-substrate friction coefficients, which are spatially modulated consistently with the localization of focal adhesions close to the cell periphery (Fig. 4a).

**Fig. 4.**
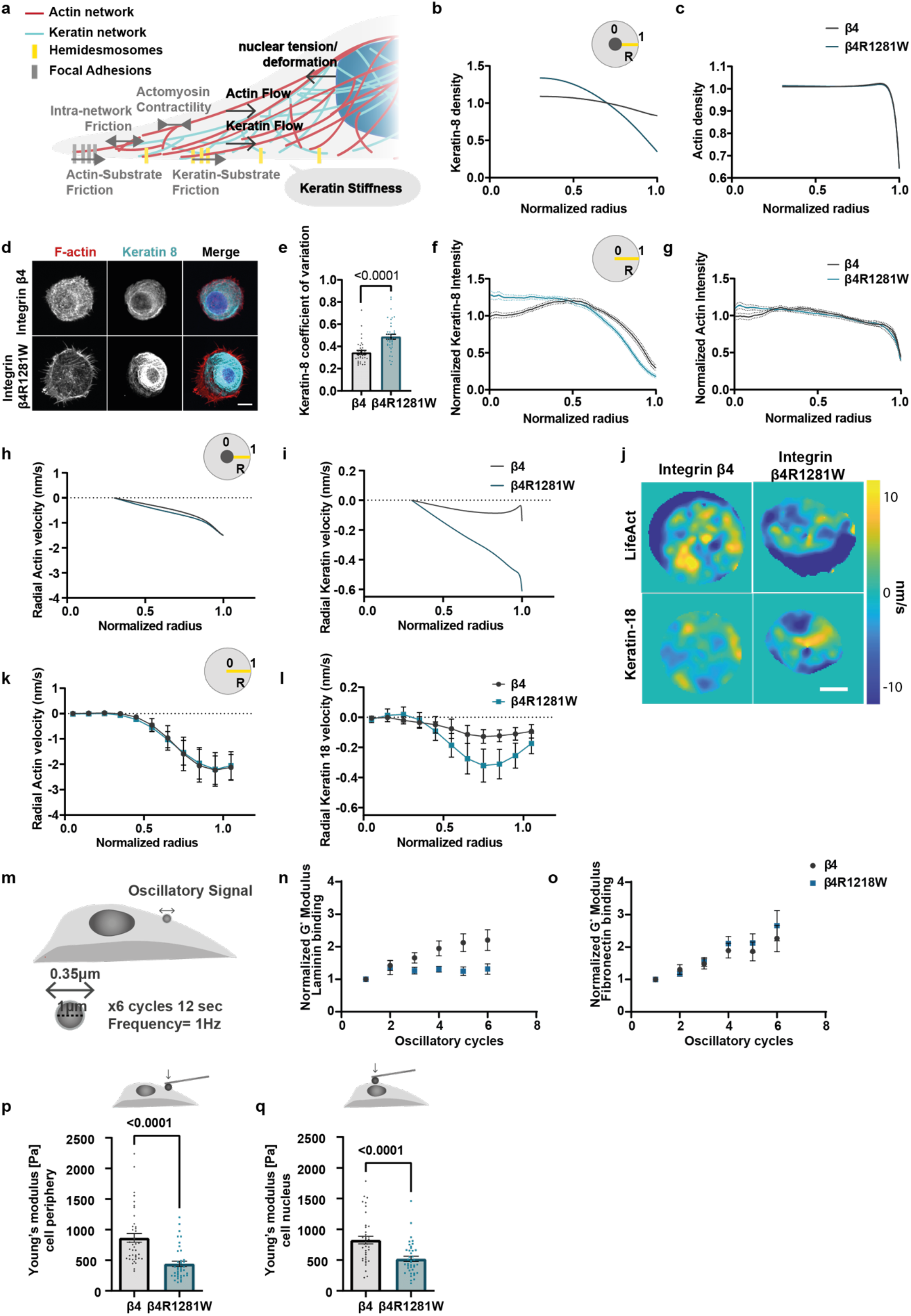
The Laminin-integrin β4-keratin link regulates cytoskeletal dynamics and cell mechanical properties. **a.** Cartoon depicting the main elements of the theoretical model. Actomyosin contractility leads to a retrograde flow of actin filaments, which is resisted by a friction between actin and the substrate, and an inter-network friction between actin and keratin. Due to applied force through actomyosin, keratin also undergoes retrograde flow, which is resisted by a friction between keratin and the substrate. The system leads to a given applied tension to the nucleus through actin-nuclear links. The mechanical connection between keratin and the substrate through integrin α6β4 is modelled by regulating keratin/substrate friction, and the stiffness of the keratin cytoskeleton. Model inputs are depicted in grey, model outputs in black. **b.** Model predictions for keratin distribution along the cell radius (R=1, cell periphery, R=0, cell centre). Note that in all model predictions in this figure, predictions start at R=0.30, where the average position of the nucleus edge is assumed to be. **c.** Model predictions for actin distribution along the cell radius (R=1, cell periphery, R=0, cell centre). **d.** Phalloidin (F-actin) and keratin 8 stainings of MCF10A cells overexpressing WT or β4R1281W integrin β4 on circular laminin patterns of 30μm diameter; scale bar is 10μm. **e.** Corresponding quantification of coefficient of variation of keratin 8 signal at the basal layer of the cell periphery (*P*<0.0001, two-tailed unpaired t-test, n= 35/40 for β4/β4R1281W from 3 independent experiments). **f.** Normalized keratin-8 intensity along the cell radius (R=1, cell periphery, R=0, cell centre) for WT (β4) or mutant integrin β4R1281W cells. (*n* = 27/43 cells for β4/β4R1281W from 3 independent experiments). The combined effect of β4 mutation and radial distribution of keratin intensity is significant (*P* < 0.0001, two-way repeated measures ANOVA). **g.** Normalized phalloidin signal intensity along the cell radius (R=1, cell periphery, R=0, cell centre (*n* = 27/43 cells for β4/β4R1281W from 3 independent experiments). The combined effect of β4 mutation and radial distribution of keratin intensity is significant (*P* < 0.0001, two-way repeated measures ANOVA). **h.** Model prediction of radial retrograde flows of actin along the cell radius (R=1, cell periphery, R=0, nucleus). **i.** Model prediction of radial retrograde flows of keratin along the cell radius (R=1, cell periphery, R=0, nucleus). **j.** Snapshot colourmaps of Life-Act-GFP (top panel) or keratin-18-mcherry (bottom panel) velocities (nm/sec) as calculated by PIV; scale bar is 10 μm. **k.** Experimental quantifications of average actin retrograde flows along the cell radius (n=12 cells for both β4 and β4R1281W). The combined effect of β4 mutation and radial actin velocities is not significant. (*P*= 0.6927, two-way repeated measures ANOVA). **l.** Experimental quantifications of keratin-18 retrograde flows along the cell radius (n=12 cells for both β4 and β4R1281W). The combined effect of β4 mutation and radial keratin velocities is significant (*P*=0.0296, two-way repeated measures ANOVA). **m.** Graphical representation of optical tweezers experimental setup. 1 μm silica beads were coated with laminin or fibronectin, and an oscillatory signal of 0.35 μm amplitude was applied with a frequency of 1 Hz upon bead attachment to integrin β4 or β4R1281W overexpressing cells. **n.** Evolution of bead resistance to force (quantified through the complex shear modulus G*) for β4 or β4R1281W overexpressing cells upon repeated cycles of oscillatory stimulations with laminin-coated beads (n=37/ 32 for β4/β4R1281W cells, mean of 3 independent experiments). The effect of β4 mutation is significant (*P* =0.0055 paired Mixed-effects model (REML)). The effect of the number of oscillatory cycles is significant. (*P* < 0.0001, paired Mixed-effects model (REML)). **o.** Evolution of G moduli for β4 or β4R1281W overexpressing cells upon repeated cycles of oscillatory stimulations with fibronectin-coated beads (n=30/ 36 for β4/β4R1281W cells, mean of 3 independent experiments). The effect of β4 mutation is not significant. (*P* =0.5988), the effect of the number of oscillatory cycles is significant (*P* < 0.0001, paired Mixed-effects model (REML)). **p.** Young’s modulus of the cell cytoplasm for β4 or β4R1281W overexpressing cells (*P* < 0.0001, unpaired two-tailed t-test, n= 40/39 for β4/β4R1281W cells, mean of 3 independent experiments). **q.** AFM stiffness measurements above the cell nucleus for β4 or β4R1281W overexpressing cells (*P* < 0.0001, unpaired two-tailed t-test, n= 38/40 for β4/β4R1281W cells, mean of 3 independent experiments). Error bars represent mean ± s.e.m.

Since integrin β4 binds to the keratin cytoskeleton and to the surrounding substrate, it can be understood as a “docking site” for the keratin network to the substrate^59^. This incorporation will have two effects. First, it will decrease the ability of actomyosin retrograde flow to drag the keratin network. We incorporate this to the model by lowering the friction coefficient between the keratin network and the substrate in the case of the integrin β4R1281W mutant, due to the absence of stable integrin β4-keratin connections (Fig. 4a). Second, the incorporation of the keratin network to hemidesmosomes will increase the crosslinking of keratin filaments^60^, which in turn will stiffen the keratin network^61,62^. Since in our model keratin-substrate friction accounts for the mechanical effect of hemidesmosomes, we included this effect by increasing the elastic modulus of the IF network proportionally to the keratin-substrate friction.

We then assessed model predictions of cytoskeletal organization and dynamics. First, the model predicts that actomyosin flows drag keratin more efficiently in mutant cells, due to decreased friction with the substrate. This will lead to keratin accumulation around the cell nucleus, whereas the organization of actin will remain largely unaffected due to its higher turnover rate (Fig. 4b,c). To validate this prediction, we seeded integrin β4 or β4R1281W expressing cells on round laminin patterns of 30μm diameter (Fig. 4d). We first assessed the homogeneity of the keratin network organization by calculating the coefficient of variation of the keratin 8 signal at the cell periphery.

This confirmed our previous observations with integrin β4 blocking antibody experiments (Fig. 3c), showing an increase for β4R1281W integrin expressing cells (Fig. 4e), which indicates a disrupted keratin organization. We then used this keratin 8 signal to calculate the distribution of the keratin network across the cell radius, where R=1 is the cell periphery as marked by actin staining and R=0 is the cell centre (Fig. 4f). The results of keratin distribution were in marked agreement with model predictions, where keratin 8 had a more central (perinuclear) localization in integrin β4R1281W expressing cells (Fig. 4f and Supplementary Fig. 4a). Accordingly, actin had a more homogeneous distribution throughout the cell in both conditions (Fig. 4g and Supplementary Fig. 4b).

A second set of predictions from the model is that both actin and keratin should exhibit retrograde flows, that keratin flows should generally be lower than actin flows, and that keratin flows should be higher in mutant integrin β4-expressing cells due to reduced friction (Fig. 4h,i). To test these predictions, we used the same circular patterns and we performed time lapse imaging of control or mutant integrin β4 cells transfected with LifeAct-GFP or keratin-18-mCherry (the binding partner of keratin-8). Then, we applied Particle Image Velocimetry (PIV) to calculate actin and keratin-18 flows (Fig. 4j and Supplementary Videos 1 and 2). Experimental values and trends for retrograde flows (measured through radial velocities) were in agreement with model predictions (Fig. 4k,l). Noticeably, overall velocities (including non-radial) were higher for the mutant condition, especially for the keratin network, indicating a looser network (Supplementary Fig. 4c,d and Supplementary Videos 1 and 2). Interestingly, the model does not predict large effects of the β4R1281W integrin mutant on traction forces, because intracellular forces are transmitted to the substrate through the actin cytoskeleton (Supplementary Fig. 4e). This is consistent with our finding that blocking or interfering with integrin β4-keratin binding did not affect traction forces (Supplementary Fig. 4f, Fig. 2e,f and Supplementary Fig. 3e,f). Thus, the link between keratin and the substrate through integrin β4 can withstand actomyosin mediated contractility, affecting the organization of the keratin network and decreasing its retrograde flow.

### A stable keratin-laminin link increases cellular stiffness

A major assumption of our model is that modulating the keratin-laminin link through β4 integrins should affect keratin crosslinking, and therefore the stiffness of the keratin network ^61,62^. The keratin network is the main cytoskeletal component contributing to the mechanical integrity of most epithelial tissues^49,63–65^. Therefore, apart from cytoskeletal organization and dynamics, the keratin-laminin linkage should also affect cellular mechanics. To show that the laminin-integrin β4-keratin link can change the mechanical resistance of the cytoskeleton, we carried out optical tweezer experiments. We coated 1 μm silica beads with laminin-111, placed them in contact with the cell surface until they attached, and then oscillated the beads horizontally by 0.35 μm at 1 Hz (Fig. 4m). Upon consecutive cycles of bead oscillations, we recorded a gradual increase in the mechanical resistance of the cytoskeleton, as quantified by the complex shear modulus G* ^66^ (Fig. 4n). Consistent with our hypothesis, this increase in the mechanical resistance was largely abrogated for cells transfected with mutant integrin β4R1281W, which has a lower affinity for plectin and therefore for keratin filaments (Fig. 4n). Confirming the specific role of laminin-integrin β4 links, the differences between WT and R1281W integrin β4 were lost when beads were coated with fibronectin instead of laminin, which binds to integrins other than β4 and can trigger actin-mediated reinforcement^55,67^ (Fig. 4o). We thus show that direct local force application to laminin-integrin β4-keratin connections increases the mechanical resistance of the cytoskeleton.

To assess not only local but also cell-level mechanical effects, we measured cell stiffness using Atomic Force Microscopy (AFM). In agreement with model assumptions, we found that cell stiffness was significantly higher in cells transfected with WT compared to R1281W integrin β4, both at the cell periphery and above the cell nucleus (Fig. 4p,q). These results indicate that a stable connection of keratins with the laminin substrate can in fact alter the mechanical properties of the cytoskeleton, eventually affecting the mechanical resistance of the cells.

### An intact keratin network protects the nucleus from actin-mediated deformation

Once the overall effects on the cytoskeleton were established, we assessed how those would affect the nucleus. We and others have previously shown that force transmission to the nucleus through the actin cytoskeleton can deform the nucleus, leading to YAP nuclear entry^3,4^. We therefore assessed if differences in mechanical deformation of the nucleus could explain the altered nuclear mechanoresponses of mutant integrin β4 cells. We first calculated model predictions of applied mechanical tension at the nucleus, which showed that the nucleus is under tension from myosin pulling forces. Due to the reduced friction with keratin, this tension is transmitted more effectively in mutant β4R1281W expressing cells, increasing by ~8% (Supplementary Fig. 5a). Assuming the same nuclear stiffness in WT β4 and β4R1281W expressing cells, this difference in tension would only result in a small nuclear deformation, as quantified by a reduction in the sphericity of the nucleus of about 3%. However, our data show decreased nuclear stiffness in mutant β4R2181W cells (Fig. 4q), attributed to lower cross-linking of the keratin network. Accounting for such reduction of nuclear stiffness, our model predicts a more pronounced increase in nuclear deformation in mutant cells, as quantified by lower sphericity (yellow bars in Fig. 5a).

**Fig. 5.**
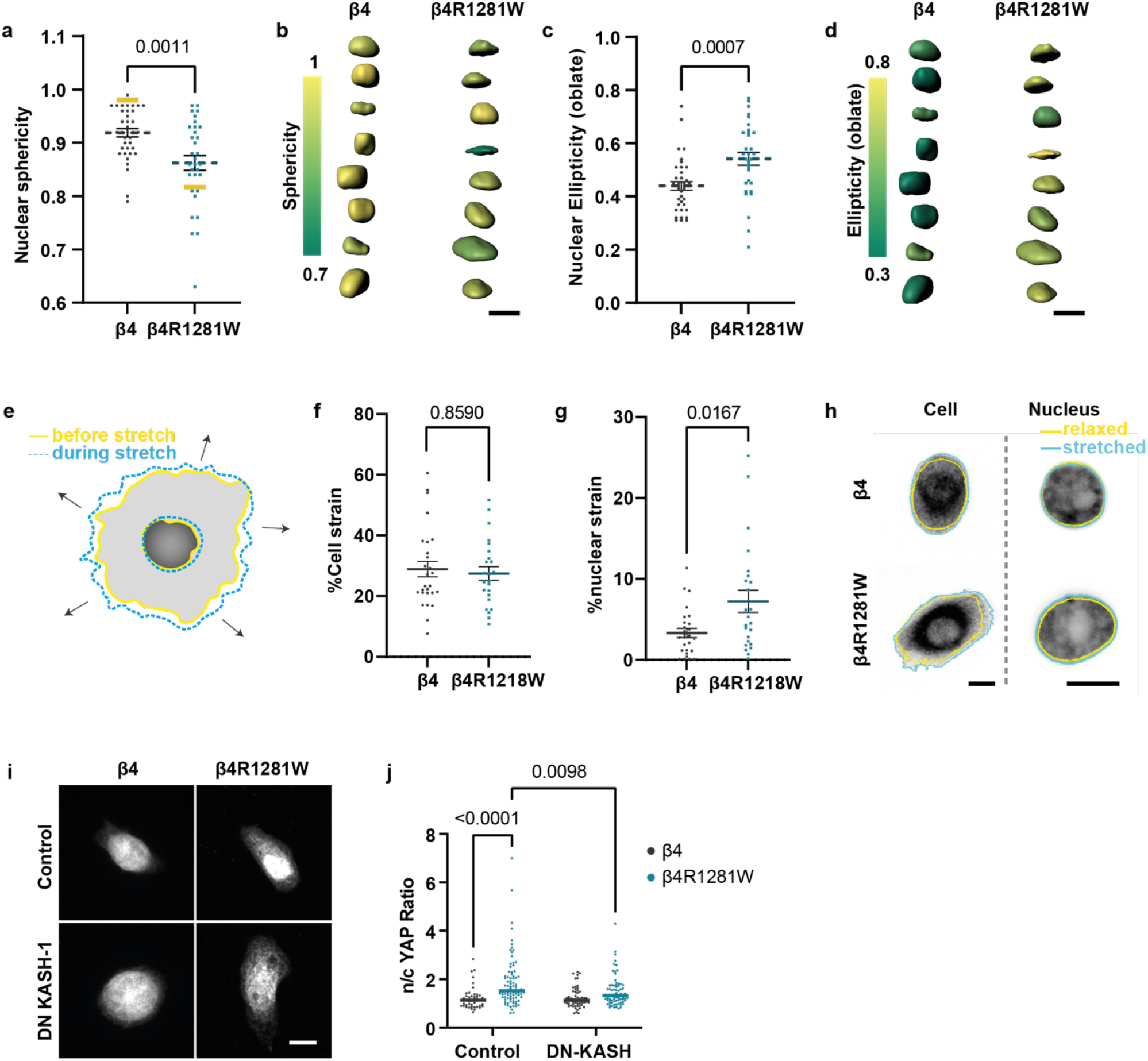
The Laminin-Integrin β4-keratin link shields the nucleus from actin-mediated deformation. **a.** Nuclear sphericity measurements (*P*=0.0011, unpaired two-tailed t-test, n=37/32 for β4/β4R1281W overexpressing cells in 3 independent experiments). Yellow bars show model predictions. **b.** 3-D segmentation and sphericity colour-coded nuclei for integrin β4 (left) and β4R1281W (right) overexpressing cells; scale bar is 10 μm. **c.** Nuclear ellipticity (oblate) measurements (*P*=0.0007, two-tailed unpaired t-test n=37/32 for β4/β4R1281W overexpressing cells in 3 independent experiments). **d.** 3-D segmentation and ellipticity (oblate) colour-coded nuclei for integrin β4 (left) and β4R1281W (right) transfected cells; scale bar is 10 μm. **e.** Illustration of cell and nuclear strain upon stretching. **f.** Cellular strain upon equiaxial stretch integrin β4 and β4R1281W overexpressing cells (*P*>0.05 two-tailed Mann-Whitney test, n=27/24 for β4/β4R1281W in 3 independent experiments). **g.** Nuclear strain for integrin β4 and β4R1281W overexpressing cells (*P* = 0.0167, two-tailed Mann-Whitney test, n=27/24 for β4/β4R1281W in 3 independent experiments). **h.** Examples of cell and nuclear strain for integrin β4 and β4R1281W overexpressing cells; scale bar is 10 μm. **i.** Control or DN-KASH transfected cells stained for YAP. Representative images of cells seeded on 11 kPa gels; scale bar is 10μm. **j.** n/c YAP ratios for control or DN-KASH transfected integrin β4 and β4R1281W cells. (Control n=48/88 and DN-KASH n= 63/76 for β4/β4R1281W in 4 independent experiments, two-way ANOVA, Bonferroni correction for multiple comparison) Error bars represent mean ± s.e.m.

To experimentally quantify nuclear deformations, we segmented 3D nuclear shapes for WT integrin β4 and β4R1281W expressing cells seeded on 11kPA PAA gels, where the differences in nuclear YAP levels were highest between the two cell lines. Indeed, nuclear shapes were markedly different between cell lines, and sphericity was lower in mutant cells (Fig. 5a,b). Whereas experimental differences were somewhat smaller than model predictions –possibly due to contributions of factors other than keratins to nuclear mechanical properties–, the trends were in agreement, and predictions fell within the experimental range of values. Overall, mutant β4R2181W cells were less spherical because they were more oblate (Fig. 5c,d), and therefore more flattened in the horizontal dimension. Further, the degree of oblate ellipticity positively correlated with the nuclear YAP intensity levels (Supplementary Fig. 5d). On the other hand, WT β4 expressing cells resembled a more prolate ellipsoid, which had an inverse correlation with nuclear YAP intensity levels (Supplementary Fig. 5b,c and e). No changes in nuclear volume were observed between the two cell types (Supplementary Fig. 5f).

Then, we explicitly evaluated whether nuclei in β4R1281W integrin transfected cells had decreased mechanical shielding. To this end, we stretched cells using a custom-built stretch device^68^ and measured the cellular and nuclear strain (Fig. 5e). Cells transfected with either WT or β4R1281W integrins stretched to the same degree (Fig. 5f,h). However, the cell nucleus stretched less in WT than in R1281W β4-transfected cells (Fig. 5g,h), confirming that the laminin-keratin link protected the nucleus from mechanical deformation. Finally, we verified whether our observed effects in YAP nuclear localization were indeed explained by changes in actin-mediated force transmission to the nucleus. We co-transfected cells with either WT or R1281W integrin β4, and DN-KASH, the KASH domain of nesprin-1. This domain binds to its binding partner in the nuclear lamina protein SUN, acting as a dominant negative that prevents binding of endogenous nesprin-1 and nesprin-2. This disrupts the linker of the nucleoskeleton and cytoskeleton (LINC) complex, preventing force transmission from the actin cytoskeleton to the nucleus ^69^. DN-KASH expression decreased YAP n/c ratios in R1281W β4-transfected cells, bringing them down to levels similar to WT-transfected cells (Fig. 5i,j), thereby confirming the role of actin-mediated force transmission to the nucleus. Our findings thus draw a mechanism where keratin-laminin links shield the nucleus from actin mediated mechanical deformation.

To complete the mechanical characterization of our system, we carried out measurements of both nuclear shape and YAP nuclear localization after inhibiting myosin contractility (with blebbistatin), actin polymerization (by blocking specifically the Arp2/3 complex using the CK666 inhibitor^70^), or both. Interestingly and as previously described^40^, blebbistatin-treated cells had high nuclear YAP levels, were more spread and had deformed nuclei with low sphericities (Fig. 6a-d). This suggests that in the absence of myosin contractility, unopposed actin polymerization spreads and flattens cells and nuclei. Indeed, perinuclear Arp2/3-driven actin polymerization has been previously shown to drive nuclear deformation^71^. Confirming this, both nuclear flattening and YAP nuclear localization were reverted by blocking actin polymerization (Fig. 6a-d). Further supporting the control of YAP by nuclear mechanics, there was a high correlation between YAP levels and nuclear shape (Fig. 6c). As the only exception to this rule and suggesting a saturation of YAP n/c ratios, blebbistatin-treated integrin β4R1281W expressing cells decreased nuclear sphericity with respect to WT integrin β4 expressing cells without further increasing YAP levels (Fig. 6c). Finally, treatment with both inhibitors diminished both nuclear YAP levels and resulted in similar levels of sphericity for both β4 and β4R1281W integrin expressing cells (Fig. 6a-d). Taken together, we show that keratin cytoskeleton stiffening can prevent actin-mediated nuclear deformation.

**Fig. 6.**
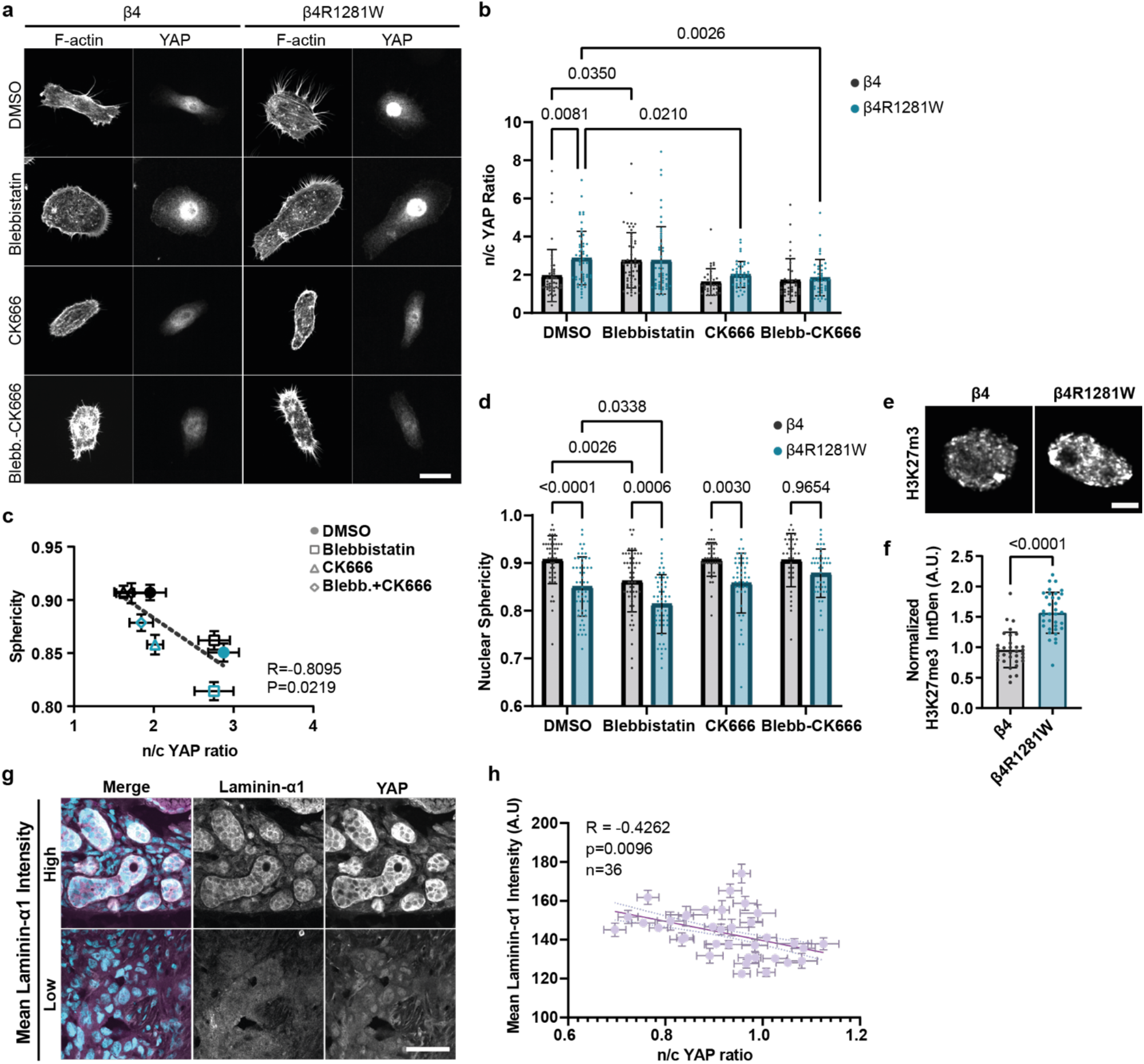
The keratin-mediated mechanoprotection is associated with nuclear shape, prevents chromatin condensation, and can also be observed *in vivo.* **a.** Phalloidin (F-actin) and YAP stainings for integrin β4 and β4R1281W overexpressing cells treated with DMSO, 25 μM Blebbistatin, 50 μM CK-666 or Blebbistatin + CK-666; Scale bar is 20 μm **b.** n/c YAP ratio quantification (n=47/53, 54/52, 37/45, 35/42 for integrin β4 and β4R1281W overexpressing cells treated with DMSO, 25 μM Blebbistatin, 50 μM CK-666 or 25 μM Blebbistatin + 50 μM CK-666, respectively, from 3 independent experiments). The effect of both β4 mutation (*P*=0.0079) and treatment (*P* < 0.0001) is significant (two-way ANOVA, Bonferroni correction for multiple comparison). **c.** n/c YAP ratios and sphericity correlation for WT β4 (black) or β4R1281W (blue) expressing cells treated with different drugs (R is Spearman’s correlation coefficient). **d.** Nuclear sphericity quantifications (n=47/53, 54/52, 37/45, 35/42 for integrin β4 and β4R1281W transfected cells treated with DMSO, 25 μM Blebbistatin, 50 μM CK-666 or 25 μM Blebbistatin + 50 μM CK-666, respectively, from 3 independent experiments). The effect of both β4 mutation and treatment is significant. (*P* < 0.0001, two-way ANOVA, Bonferroni correction for multiple comparison). **e.** Nuclei stained for H3K27me3 stainings; scale bar is 5 μm. **f.** Quantification of H3K27me3 normalized Integrated Density (*P*<0.0001 two-tailed unpaired t-test, n=33/35 for β4/β4R1281W from 3 independent experiments). **g.** Example of laminin-α1 stainings and YAP stainings on human breast cancer sample biopsies; scale bar is 50 μm. **h.** Correlation of laminin-α1 signal and n/c YAP ratios in human breast sample biopsies (n=36 images form 5 tumours and one control sample; R is Spearman’s correlation coefficient). Error bars represent mean ± s.e.m.

### Laminin-Keratin mediated nuclear shielding prevents chromatin condensation and can also be observed *in vivo*

Apart from nuclear YAP translocation, forces applied to the nucleus can trigger different mechanotransduction events, which could also take place in our system. For instance, Lamin A/C levels have been shown to affect nuclear mechanics and nuclear shape integrity^72,73^, and nuclear deformation regulates Lamin A/C levels^74^. We found no significant difference between WT and β4R1281W cells in the overall expression levels of lamin A/C, indicating that the nuclear lamina was not compromised (Supplementary Fig. 6a-c).

Apart from the nuclear lamina, there is a growing evidence of the emerging role of chromatin as a dynamic responsive element to external mechanical inputs^75^. It has been shown that upon sustained mechanical strain there is an increase in H3K27me3, a histone mark indicative of compacted heterochromatin, influencing the capacity for cell differentiation^76,77^. More recently, strain-induced changes in heterochromatin organization and upregulation of H3K27me3 were accompanied by changes in gene expression related to hemidesmosome assembly^77^. We thus stained WT and mutant β4 cells with H3K27me3 to evaluate the levels of chromatin condensation. We found that cells expressing mutant integrin β4R1281W had higher levels of this heterochromatin mark as compared to WT expressing cells (Fig. 6e-f). Thus, laminin-keratin links shield the nucleus mechanically and impair not only YAP signaling, but also other effects of nuclear mechanotransduction.

Finally, we assessed whether the role of laminin-111 in dampening nuclear mechanoresponses was also observed *in vivo*. To this end, we quantified laminin-α1 and n/c YAP ratios in healthy human mammary glands or invasive ductal carcinomas of no specific type (IDC-NST). Supporting the physiological relevance of our findings, we found that there was an overall negative correlation between the laminin-α1 levels expressed by cancer cells and n/c YAP ratios (Fig. 6g,h).

Our findings draw a novel picture on how ECM can modulate cellular responses and can have important implications not only during disease progression where BM integrity is compromised, but also during cell differentiation and tissue morphogenesis.

## Discussion

Our results demonstrate a context-specific mechanoresponse, where laminin-111 impairs cellular response to substrate rigidity. This could help interpret several previous results in the literature. Loss of laminin-111 expression has been correlated with breast cancer progression^22^. However, in Ductal Carcinoma in Situ (DCIS) where laminin levels may still be uncompromised, YAP was not nuclear despite changes in the rigidity of the tissue, and nuclei appeared smaller and rounder compared to invasive ductal carcinoma (IDC) models that expressed a subset of YAP target genes^11^. Moreover, increased binding of integrin α6β4 to basement membrane-rich alginate networks prevented the stiffness-induced malignant phenotype of mammary epithelial cells^7^. These findings are in alignment with earlier studies in an *in vitro* breast cancer model demonstrating a role for integrin α6β4 in maintaining 3D cell organization and polarity, and blocking integrin β4 triggered a malignant phenotype^23^. Finally, features such as cell softening in the invasive front of mammary epithelial spheroids have been associated with both changes in cell and nuclear shapes, and a differential expression of keratins, supporting an association between the keratin cytoskeleton and cell stiffness and invasiveness^78–80^. Indeed, changes in keratin expression during breast cancer invasion (i.e., from a luminal keratin 8 to a basal keratin 14)^79^ could also affect the mechanical properties and responsiveness of the keratin network to external signals. We thus propose that cell engagement to laminin, and its subsequent effect on nuclear and cell mechanics through the keratin cytoskeleton, could affect tissue organization and cell responses. Loss of these interactions could promote a malignant phenotype by rendering the nuclei more susceptible to deformation. These context-specific nuclear mechanoresponses could possibly explain the often contradicting roles of YAP in breast cancer progression^11,81–83^. Similarly, α6β4 integrins have been reported to both induce and inhibit malignant phenotypes^37,84^. The tumor-promoting role of α6β4 involves contexts likely unrelated to mechanical effects on keratins, as it is maintained for cells in suspension^24^ or lacking the extracellular domain of α6β4^85,86^, or involves a switch of α6β4 from keratin to actin binding^87^.

Intermediate filaments (IFs) are largely considered as the stronghold of cell and tissue integrity. Recently, both keratin and vimentin networks have been shown to provide mechanical support to the cell nucleus^46,88,89^. This mechanical role of IFs is known to be regulated by several biochemical intracellular signals, such as phosphorylation or divalent cations^90–92^. Through this work, we show that the mechanical protection exerted by IFs can also be tuned by external signals such as ECM composition and stiffness, and that this can have implications in chromatin compaction. Thus, changes in the ECM could eventually lead to altered gene expression and differentiation. For instance, repeated cycles of mechanical strain in epidermal progenitor cells (EPCs) led to a decrease in nuclear actin levels and a force driven perinuclear actin polymerization, resulting in H3K27me3-mediated gene silencing, eventually attenuating lineage commitment^76^. Responses of this kind may be modulated through a mechanical protection by keratins. This could be relevant in the many scenarios where laminin and keratin play a role, from cancer to very early stages of embryonic development, and cell differentiation events^93–95^.

## Acknowledgments

We thank A. Farré and other members of IMPETUX OPTICS, S.L for their help and expertise in the design and implementation of the optical tweezers experiments, R. Sunyer for help and advice with microprinting experiments, S. Usieto, A. Menéndez, N. Castro and W. Haaksma for providing technical support, L. Rosetti and S. Saloustros for providing data analysis tools, and J. de Rooij, A.L. Le Roux, L. Faure, A. Labernadie, R. Oria, A. Elósegui-Artola, and all the members of P. Roca-Cusachs and X. Trepat groups for helpful discussion. Finally, we thank G. Wiche, A. Sonnenberg and N. Montserrat for providing plasmids, antibodies or cell lines used for this work. We acknowledge funding from the Spanish Ministry of Science and Innovation (PGC2018-099645-B-I00 to X.T., PID2019-110949GB-I00 to M.A. and P.S., BFU2016-79916-P and PID2019-110298GB-I00 to P.R.-C.), the European Commission (H2020-FETPROACT-01-2016-731957), the European Research Council (CoG-616480 to X.T., CoG-681434 to M.A), the Generalitat de Catalunya (2017-SGR-1602 to X.T. and P.R.-C., 2017-SGR-1278 to M.A. and P.S.), European Union’s Horizon 2020 research and innovation programme under the Marie Skłodowska-Curie grant agreement No 797621 to M.G.-G. The prize “ICREA Academia” for excellence in research to M.A. and P.R.-C., Fundació la Marató de TV3 (201936-30-31 and 201903-30-31-32), and “la Caixa” Foundation (Agreement LCF/PR/HR20/52400004). IBEC and CIMNE are recipients of a Severo Ochoa Award of Excellence from MINCIN.

## Author contributions

Z.K. and P.R.-C. conceived and supervised the project. Z.K., M.Z., I.A., T.K and A.E.M.B. performed experiments. M.A. and P.S conceived the computational model. P.S. implemented the computational model and performed simulations. M.G.-G. developed software and analysed data. P.W.B.D., X.T. and M.A. contributed to technical expertise, materials and discussion. Z.K. and P.R.-C. wrote the manuscript. Z.K., P.S., A.E.M.B, P.W.D.D., X.T., M.A., P.R.-C edited and reviewed the manuscript.

## Declaration of interests

The authors declare no competing interests.

## Methods

### Cell culture

Mammary epithelial cells (MCF 10A) were purchased from ATCC. Cells were used for maximum 18 passages and were cultured in DMEM-F12 (LifeTechnologies, 21331-020) with 5% horse serum, 1% penicillin streptomycin, EGF (20 ng ml−1), hydrocortisone (0.5 μg ml−1), cholera toxin (100 ng ml−1), and insulin (10 μg ml−1). Human breast myoepithelial immortalized cell lines have been described previously^96,97^ and used for maximum 8 passages. They were cultured in Hams-F12 (Sigma, N4888) media supplemented with 10% foetal bovine serum (FBS), 1% penicillin streptomycin, hydrocortisone (1 μg ml−1), epidermal growth factor (EGF; 10 ng ml^−1^) and insulin (5 μg ml^−1^). All cells were regularly tested for mycoplasma contamination.

### Preparation of polyacrylamide gels

Polyacrylamide gels were prepared as described previously ^97^. Briefly, glass bottom MatTek dishes and slides were activated with a solution of acetic acid, 3-(Trimethoxysilyl)propyl methacrylate (Sigma), and 96% ethanol (1:1:14) for more than 10 min and 2 h for glass bottom MatTek dishes and glass slides respectively. The glass was then washed with 96% ethanol and air-dried. Different concentrations of acrylamide and bis-acrylamide were mixed to produce gels of different rigidity ^5^ and mixed together with 2% v/v fluorescent carboxylated 200 nm beads (Invitrogen), 0.05% APS (A3678, Sigma), and 0.05% v/v tetramethylethylenedi-amine (T9281, Sigma). The solution was placed on the glass and covered with a coverslip, letting the gel to set for 50 min. The coverslip was then removed, and gels were coated with 50 μl of 10% 0.5M HEPES pH 6.0, 0.004% Bis-acrylamide, 0.05% Igracure 2959 and 4% of 10 mg/ml Acrylic-acid NHS/DMSO (A8060, Sigma) in milliQ water. The gels were then exposed to UV light (XX-15, UVP) 365nm wavelength for 5min, washed twice with a 0.5M HEPES pH 6.0 solution followed by 2x PBS solution washes. Gels were then incubated overnight at 4 ^o^C with 10μg/ml solution of laminin-111 (L2020, Sigma), collagen I (First Link, UK) or fibronectin (F0895, Sigma-Aldrich) protein solution in PBS. The rigidity of polyacrylamide gels was measured and characterized with Atomic Force Microscopy as described previously ^97^.

### Immunostaining

For immunostaining, cells were fixed with 4% paraformaldehyde for 10 min, permeabilized with 0.1%Triton X-100 for 4 min, after a 30min blocking step with 0.5% fish gelatin, cells were incubated with primary antibodies (1.5h, room temperature in 0.5% fish gelatin in PBS), followed by incubation with secondary antibodies. When Phalloidin -Atto 488, (Sigma-Aldrich, Cat# 49409), - TRITC, (Sigma-Aldrich, Cat# P1951), -iFluor 647 Reagent (Abcam, Cat# ab176759) was used it was added with the secondary antibodies. Hoechst 33258 staining dye was used for nuclear labelling following a 10min incubation at room temperature.

The primary antibodies used, and their respective dilutions are: Rabbit Phospho-Paxillin 1:100 (Tyr118) (Cell Signaling, Cat# 69363 and 2541s), rabbit anti-YAP (D8H1X) XP® 1:100 (Cell Signaling, Cat# 14074), mouse anti-YAP1 YAP1 (63.7) 1:100 (Santa Cruz, Cat# sc-101199), rabbit anti-Cytokeratin 8 [EP1628Y] 1:200 (Abcam, Cat# ab53280), rabbit anti-plectin antiserum 1:400 (#46, gift from Gerhard Wiche^98^), mouse anti-Integrin β4, clone ASC-3, 1:100 (Merck Life Science, S.L., Cat# MAB2058), mouse Anti-Lamin A + Lamin C antibody [131C3] 1:200 (Abcam, Cat# ab8984), mouse anti-Lamin A/C (E1) 1:100 (Santa Cruz, Cat# sc376248), rabbit anti-Tri-Methyl-Histone H3 (Lys27) (C36B11) 1:300 (Cell Signaling, Cat# 9733).

The secondary antibodies used are: mouse Alexa Fluor −488 (Cat# A-11029), −555 (Cat# A-21424),-647 (Cat# A-21236) and rabbit Alexa Fluor −488 (Cat# A-21206), −555 (Cat# A-21429), −647 (Cat# A-21245) all at 1:300 concentration (ThermoFisher).

### Image acquisition

Immunofluorescence images and actin/keratin retrograde flow experiments were taken in a Nikon TiE inverted microscope with a spinning disk confocal unit (CSU-WD, Yokogawa) and a Zyla sCMOS camera (Andor), using 60x objective (plan apo; NA, 1.2; water immersion). Epifluorescence images were taken on an automated inverted microscope (Nikon Eclipse Ti) using MetaMorph/NIS Elements imaging software and 60x objective (plan apo VC; NA, 1.4; oil immersion).

### Immunostaining analysis

Fiji software was used to perform the image analysis ^99^. The length of p-Paxillin focal adhesions was assessed as described previously^100^ using maximum projection of confocal images, by measuring the length of bright focal adhesions (FA) on the edge of single cells and averaging the length of 10 FAs per cell. YAP n/c ratios were calculated as described previously^3^ using average projections of confocal images and by dividing the intensity on a nuclear region and a region with equal size in the cytosol immediately adjacent to the nuclear region upon correcting for background. The corresponding Hoechst staining image and fluorescent staining signals were used to delimit nuclear versus cytosolic regions. For Supplementary Fig.1 and 3e-g epifluorescence images were also used instead of confocal stacks. Due to the large number of conditions, each experimental repeat of FA and YAP measurements in Fig. 2a,c included the control condition, but could not include all conditions at the same time. To account for this, values were corrected for each stiffness by the fold-difference between the control value of the experiment, and the average value of all control samples. Quantification of the coefficient of variation of the keratin-8 signal was carried out by measuring the standard deviation of the signal in 3 areas around the cell periphery marked by the actin signal at the basal surface of single cells and averaged for each cell. The standard deviation was then normalized to the corrected mean intensity of the fluorescent signal. Corrected integrated density of keratin-8 (Supplementary Fig. 3a) and lamin A/C (Supplementary Fig. 6b) labelled cells was quantified on sum projections of single cells or nuclei and was carried out using ImageJ software. SUM H3K27m3 signal was quantified using Imaris software (© Oxford Instruments) upon 3D segmentation of cell nuclei. For each experiment SUM (integrated density) signal was normalized to the average signal of β4 to account for experimental variation.

### Western blots

Western blots were implemented following standard procedures. Briefly, cells were lysed using RIPA buffer. Following denaturation, lysates were loaded on 4%–20% polyacrylamide gels (Bio-Rad) and transferred onto a nitrocellulose membrane (Whatman, GE Healthcare Life Sciences). After blocking, the membranes were incubated with primary antibody overnight at 4 °C and with the horseradish-peroxidase (HRP)-conjugated secondary (1:5000) (merckmillipore) antibody for 2 hr at room temperature. ECL Western Blotting Substrate (Pierce, ThermoFisher) was used to detect HRP and the bands were visualized with the ImageQuant LAS 4000 imaging system (GE Healthcare Life Sciences). The intensity of the bands was analysed using ImageJ software.

Antibodies used are: mouse anti-Lamin A/C (E1) 1:1000 (Santa Cruz, Cat# sc376248), mouse anti-Integrin beta 4 [M126] 1:1000 (Abcam, Cat# ab29042), rabbit anti-GAPDH (D16H11) XP® 1:1000 (Cell Signaling, Cat# 5174), mouse anti-GAPDH (6C5) 1:3000 (Santa Cruz, Cat# sc-32233), mouse Anti-Nesprin3 antibody [Nsp3] 1:500 (Abcam, Cat# ab123031), rabbit Phospho-Myosin Light Chain 2 (Thr18/Ser19) 1:500 (Cell Signaling, Cat# 3674), rabbit anti-Cytokeratin 8 [EP1628Y] 1:2000 (Abcam, Cat# ab53280).

### Preparation, procedure and quantification of stretch experiments

Stretch experiments were carried out using a stretch device coupled to an upright Nikon eclipse Ni-U microscope as described before^3,68^. Briefly, stretchable membranes were prepared by mixing PDMS base and crosslinker at a 10:1 ratio, spinning the mixture for 1 min at 500 rpm, and finally cured overnight at 65°C. Once cured, PDMS membranes were placed on stretching devices and coated with 10 μg/ml laminin overnight at 4°C. Cells were then seeded on the membranes and stretch experiments were caried out 4-8h post-seeding, in an upright microscope (Nikon eclipse Ni-U). Calibration of the system was done to adjust the vacuum applied to obtain 5% stretch of the PDMS surface. During stretching cells were kept in CO_2_-independent medium (ThermoFisher, Cat# 18045088) containing 1:100,000 Hoechst 33258 staining dye and 10 μg/mL of rutin (Sigma R5143) to prevent photobleaching^101^ and treated with CellTracker (Invitrogen, Cat# C34565). Membranes were subjected to equiaxial stretch upon the application of vacuum and images were acquired before and during stretch with a water immersion 60x objective (NA = 1.0). Changes in cell and nuclear strain were calculated by tracing cell and nuclear perimeter using fluorescence signal and/or bright field images and Hoechst signal respectively, before and during stretch. A value of 0.1 has been assigned to negative strain values.

### Atomic Force Experiments and quantification

Stiffness of cell nucleus and cytoplasm was measured with atomic force microscopy (AFM) using a Nanowizard 4 AFM (JPK) mounted on top of a Nikon Ti Eclipse microscope^97^. The spring constant of the cantilevers was calibrated by thermal tuning using the simple harmonic oscillator model. The Hertz model was fitted to approach curves to obtain the stiffness value using the “JPKSPM data processing” software (JPK). Cells were seeded on laminin-111-coated coverslips and a force curve on top of the nucleus and cytoplasm was acquired for each of the cells. Cells were kept at 37 °C using a BioCell (JPK) and maintained in CO_2_-independent medium (ThermoFisher Cat# 18045088). Values over 3-fold higher than the mean were excluded. In total 3 out of 180 values were excluded (2 values, β4 panel 4q, 1 value, β4R1281W panel 4p).

### Optical tweezers experiments and quantification

Optical tweezers experiments were performed using SENSOCELL device incorporated in a Ti Eclipse Nikon microscope, using a 60x objective (plan apo; NA, 1.2; water immersion). 1 μm carboxylate beads (01-02-103, micromer) were coated with biotinylated laminin (LMN03, Cytoskeleton, Inc.) or FN7-10 (a four-domain segment of fibronectin responsible for cell binding and containing the RGD and PHSRN motifs^102^) and biotinylated bovine serum albumin at a ratio of 1:10. Cells were seeded on #1.5 cover slips (Menzel-Gläser) previously coated with laminin. During the experiment cells were kept in serum-free CO_2_-independent medium containing 1% penicillin streptomycin (gibco), 1.5% 1M HEPES pH7.5 (sigma), 2% L-Glutamine (gibco). Beads were flowed into the medium and were put into contact with cells upon trapping them. A series of 6 oscillatory cycles of 12 sec each and with an amplitude of 0.35μm and frequency 1 Hz were performed with an interval of 10 sec for bead repositioning. The complex shear modulus G* was measured for each oscillation cycle using the Micro-rheology routine of the LightACE, the control software of the optical tweezers instrument. For the computation of G*, the force was determined by means of the calibration-free "momentum method" ^66^ and the particle position was obtained using the measured stiffness of the trap.

### Cell monolayer experiments

Cells were seeded on 11 kPa gels of 18 mm diameter prepared as described before on MatTek dishes. Upon functionalization, 6×9 mm magnetic PDMS gaskets were placed on top and the plates were attached to a magnetic holder. 1.2 × 10^5^ cells were seeded in the gasket and non-attached cells were washed after 5 hours. Cells were kept overnight in serum free medium prior to the removal of the magnetic gasket. For antibody blocking experiments a constant concentration (10 μg/ml) of blocking antibodies or IgG control was maintained throughout the course of the experiment.

### Micropatterning

Circular (30 μm diameter) patterns were generated using the PRIMO micropatterning platform (Alvéole) on the surface of PDMS gels made as described elsewhere^103^. Briefly, PDMS CY52-276 A and B (Dowsil) were mixed at a 9:10 ratio on ice. The solution was then used to coat 35 mm MatTek dishes and cured at 65ºC overnight. Patterns were generated as per manufacturers instruction. Beads were attached on the surface of the PDMS using an APTES-Ethanol solution (5% v/v) (Sigma-Aldrich) to identify the surface of the gel. Gels were then passivated with a 2-stepPoly-L-Lysine (PLL) (Sigma-Aldrich - P2636) - PEG-SVA (Laysan Bio - MPEG-SVA-5000) incubation. Prior to micropatterning gels were covered with (p-Benzoylbenzyl) trimethylammonium chloride (PLPP) (BocSciences) that allows UV light-induced PEG degradation. Gels were then incubated with a 10 μg/ml laminin solution (1:1 rhodamine labeled (LMN01-A, Cytoskeleton, Inc.) to non-labeled laminin-1 (L2020, Sigma Aldrich) or 1:10 labeled fibrinogen (Invitrogen) to laminin-1 (L2020, Sigma Aldrich) solution.

### Traction force microscopy

Traction force experiments were performed as described before^97^. Briefly, cells were seeded on PAA gels of different rigidities, fabricated as described above. Traction force experiments were carried out using multidimensional acquisition routines on an automated inverted microscope (Nikon Eclipse Ti) equipped with thermal, CO_2_ and humidity control using MetaMorph/NIS Elements imaging software. Fluorescent images of the beads and phase contrast images of the cells were acquired every 10min during the course of the experiment. Local gel deformation between any experimental time points and a reference image obtained after cell trypsinization were computed with a home-made particle imaging velocimetry (PIV) software (Trepat et al., 2009) in Matlab (MathWorks Inc.). Traction forces were computed using Fourier traction microscopy with a finite gel thickness^104^ and averaged for each cell.

### Actin and keratin retrograde flow experimental design and quantification

Cells were transfected with LifeAct-GFP and keratin-18-mCherry and seeded on PRIMO micropatterned PDMS, as described previously. Images were taken every 4 sec in a Nikon TiE inverted microscope with a spinning disk confocal unit (CSU-WD, Yokogawa) and a Zyla sCMOS camera (Andor) controlled by Micro-Manager^105,106^ using ×60 objective (plan apo; NA, 1.2; water immersion). The local velocity fields of the actin and keratin fluorescence signals were measured by comparing each frame and its previous timepoint with a custom-made PIV software in Matlab. A mask of each cell was drawn with respect to the F-Actin signal in ImageJ^107^. A radial coordinate, centred in the mask centroid, was assigned to each PIV data point, and normalized by the local radius of the cell mask contour. Likewise, the local velocity fields were decomposed into their radial and tangential components. The distributions of total and radial velocities inside each cell were then binned into equal sized intervals of the normalized radial coordinate. The average total and radial velocity for each radial bin was then calculated.

### Cell transfection

Non-viral cell transfection was carried out using Lipofectamine™ 3000 Transfection Reagent (Invitrogen) following manufacturer’s instructions. For the DN-KASH1 experiments, FACS selection was carried out upon transfection. EGFP-Nesprin1-KASH or mCherry-Nesprin1-KASH and EGFP empty vector control and LifeAct-GFP were described previously^3,100,108^, pcDNA3.1-mCherry (Plasmid #128744) and mCherry-Keratin-17 (Plasmid #55065) were purchased from addgene. siRNA transfection was performed using Lipofectamine™ RNAiMAX Transfection Reagent (Invitrogen), following manufacturer’s instructions. Nesprin-3 siRNA (Cat# LQ-016637-12-0002) and NTC control (Cat# D-001220-01-05) was purchased from Dharmacon™. Retroviral particles for the generation of stable integrin β4-GFP and β4R1281W-GFP (LZRS-IRES-zeo plasmids were a gift from A. Sonnenberg^39^) lines were generated in HEK293T cells expressing retroviral packaging plasmids (gift from N. Montserrat) and transfected using Lipofectamine™ 3000 Transfection Reagent (Invitrogen).

### Blocking antibody and drug treatment experiments

Antibody blocking experiments were performed by incubating the cells with Anti-Integrin ß4 Antibody, clone ASC-3 (MAB2058Z) and clone ASC-8 (MAB2059Z), Anti-Integrin α3, clone P1B5 (MAB1952Z), Anti-Integrin β1 Antibody, clone P5D2 (MAB1959Z) (Merck Life Science, S.L.) and Anti-Integrin alpha 6 antibody, clone GoH3 (ab105669) (Abcam), for 20 min at room temperature prior to cell seeding and maintained in the same concentration (10 μg/ml) of blocking antibodies throughout the duration of the experiment.

Drug treatment experiments were carried out by incubating cells with 25 μM Blebbistatin, 50 μM CK-666 or the highest corresponding concentration of DMSO (Sigma-Aldrich) for 2h upon cell attachment and spreading on 11 kPa PAA gels coated with laminin-111 (L2020, Sigma-Aldrich).

### Actin anisotropy quantification

Actin anisotropy was quantified in maximum projection images from confocal stacks labelled with phalloidin. The anisotropy quantification was implemented using the imageJ fibril tool plugin^109^.

### Keratin distribution quantification

Keratin and actin distributions were quantified on keratin 8 and phalloidin actin average projections of cells cultured in 30 μm patterns with a custom-made MATLAB code. As described above, a mask of each cell was drawn by thresholding for the actin signal using ImageJ software^107^ and a normalized radial coordinate was assigned to each point of the image. An intensity profile was calculated by binning the normalized intensity values into equal sized intervals of the normalized radial coordinate and averaging the values of intensity for each bin. Then, profiles of intensity with respect to radial distance were normalized by the integral of the curve.

### Nuclear shape quantification

3D segmentation and shape characterization of Hoechst 33258 labelled nuclei was implemented using Imaris software (© Oxford Instruments) surface segmentation. Parameters used are: sphericity, defined as: 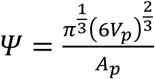 where *V_p_* is the volume of the particle and *A_p_* is the surface area of the particle; ellipsoid prolate, defined as: 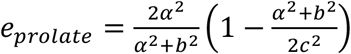 and ellipsoid oblate, defined as: 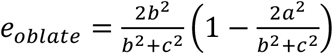, where a, b, and c are the lengths of three semi-axes determining the shape of an ellipsoid. For more information: https://www.bitplane.com/download/manuals/ReferenceManual9_2_0.pdf

### Immunofluorescence on patient tissue samples and quantification

Whole breast tissue sections diagnosed as high grade (3) invasive ductal carcinoma of no specific type (IDC-NST) was approved by the Tissue Science Committee of the University Medical Center Utrecht. Immunofluorescence was done on 4 μm whole breast tissue sections. Formalin fixed paraffin embedded IDC-NST tissue sections were deparaffinized, followed by antigen retrieval by boiling in Tris-EDTA buffer for 20 min. Sections were washed with 1x PBS and blocked with 1% BSA in PBS for 30 min before antibody and DAPI incubations. Slides were mounted in ProLong™ Diamond Antifade (Thermo Fisher P36961) and imaged after a 24h drying period.

Antibodies used were: mouse anti-Laminin α-1 (CL3087) (Invitrogene, Cat# MA5-31381), rabbit anti-YAP (D8H1X) XP® (Cell signaling, Cat# 14074). Image acquisition was performed in a Nikon TiE inverted microscope with a spinning disk confocal unit (CSU-WD, Yokogawa) and a Zyla sCMOS camera (Andor), using a 40× objective (Plan Fluor, NA 0.75, Dry). Quantification was performed in confocal slides of 5 different tumours and one healthy control. For each tumour/control sample, 2-7 images of areas defined by tumour boundaries were analysed. For each image, an average of laminin intensity was calculated, and average n/c YAP ratios were obtained by dividing the intensity on a nuclear region and a region with equal size in the cytosol immediately adjacent to the nuclear region.

### Material and software availability

The codes of this study are available from the corresponding authors on request. The computational model of actomyosin and keratin networks is available at https://gitlab.com/PSaez83/actinkeratincell2022.git. This study did not generate new unique reagents.

### Statistical analysis

Statistical analyses were performed using GraphPad Prism software (GraphPad, version 9). Statistical significance was determined by the specific tests indicated in the corresponding figure legends. Non-parametric tests were performed when both, original and log-10 transformed datasets were not normally distributed where appropriate. All experiments presented in the manuscript were repeated at least in 3 independent experiments, except: Fibronectin and collagen conditions for Supplementary Fig. 1 (n=2), Supplementary Fig. 3 h,j (n=2), Supplementary Fig. 6c (n=2).

### Computational model of actomyosin and keratin networks

#### Geometric model and model considerations

To understand the mechanics of the actomyosin and keratin networks of the cell and the mechanical interaction between these two networks and between the networks and the substrate, we developed a mathematical model of the cell composite cytoskeleton, which we solved numerically using the finite element method. We consider a 1D domain of length L modelling the region comprised between the nucleus edge and the cell edge, for a stationary and radially symmetric cell (Method Fig. 1a). Our model is composed of an F-actin network that is actively pulled by myosin motors, as modelled previously^57,110,111^, and a viscoelastic network of intermediate filaments (IF) that is dragged by the F-actin flow, as modelled previously^112^. We show a sketch of the model in Method Fig. 1b. We thus consider a cellular domain along coordinate x and given by (0, L), where x = 0 represents the nuclear boundary of the composite network and x = L represents the cell front (Method Fig. 1a). We assume that each network has a well-defined hydrodynamic velocity field, *v^a^*(x) for actin and *v*^*IF*^(x) for IF, and can sustain a mechanical stress, *σ^a^*(x) and *σ^IF^*(x) respectively. Furthermore, these networks interact with their surroundings and between each other frictionally, modeling both unspecific mechanisms and the net mechanical effect of transient binding between specific ligands. Each network possesses distinct constitutive laws as detailed next.

**Method Fig. 1:**
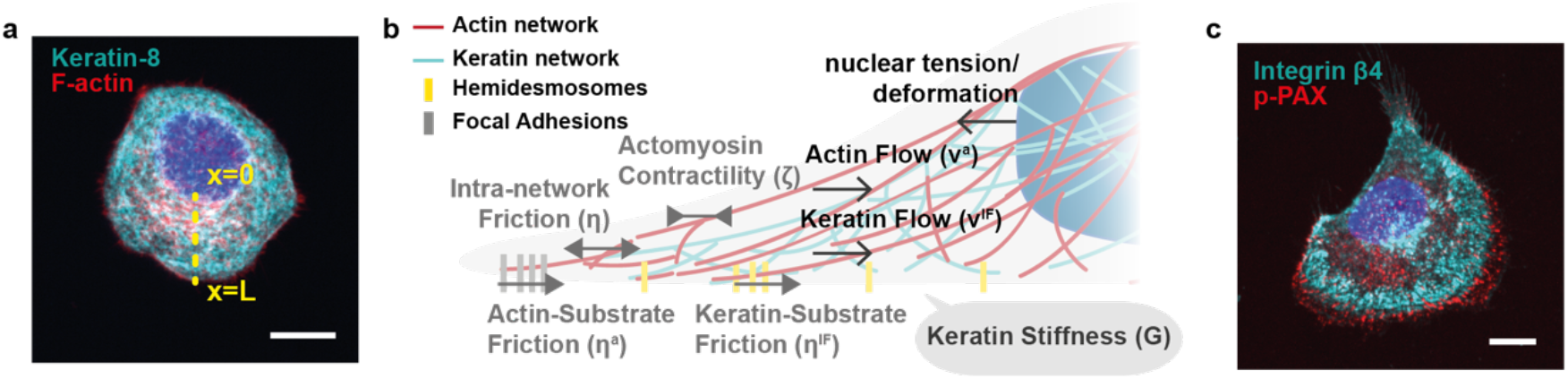
(a) Representation of cellular structure depicted by the model. A half 1D representation of the cell is drawn, with x=L and x=0 representing the front and nuclear boundaries of the cell; scale bar is 10μm. (b) Geometry and intracellular structure of the model. The network of actin fibres is shown in red and keratin fibres in cyan. The extranuclear networks can impose compressive and tensional forces on the nucleus, leading to nuclear deformation. (c) Focal adhesions, as depicted by phospho-Paxillin staining (pPAX in red) and hemidesmosomes (integrin β4 adhesions in cyan) are spatially accumulated in the periphery of the cell; scale bar is 10μm.

#### Mechanics of the F-Actin network

To model the mechanical behaviour of the F-Actin network and, ultimately, compute the velocity of the F-actin network, *v^a^*, we describe the balance of linear momentum neglecting inertial forces along with boundary conditions as described previously^57,110,111^:

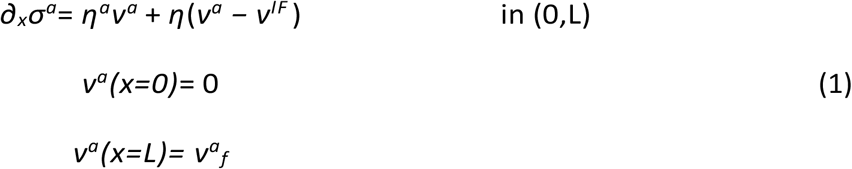

where we considered *v*^*a*^_*f*_ = −1.5×10^−3^ μm/s for the actin velocity at the cell edge, modelling polymerization at the lamellipodium, and zero actin velocity at the nucleus, in agreement with experimental results (main text Fig. 4k). The second term in the right-hand side corresponds to frictional forces due to the relative motion between actin and the IF network controlled by the constant friction coefficient *η*. The first term models the frictional forces resulting from sliding of the actin network relative to the extracellular matrix (ECM) with friction coefficient *η^a^*. We modulate friction between actin and the ECM, *η^a^*, in space following a sigmoidal distribution:

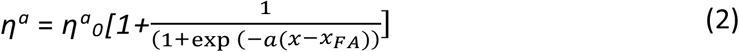

Which adopts a baseline value *η*^*a*^_*0*_ close to the nucleus, increases to nearly 2*η*^*a*^_*0*_ at the cell front, with *η*^*a*^_*(xFA)*_=0.5*η*^*a*^_*0*_ for *xFA* = 0.8*L*. *α* is a constant that sets the slope of the sigmoidal function. This space-dependent function accounts for the fact that focal adhesions mainly form at the periphery of the cell^113,114^, as we also show in Method Fig. 1c. This parameter is important because we model mutant cells to have a weaker interaction with the laminin ECM, by reducing *η*^a^_0_.

We consider the constitutive relation for the internal stresses of the F-actin network as:

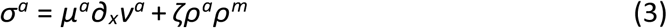

where the first term considers the viscosity of the actin network and the second term models the contractile forces of the myosin motors. *μ^a^* is the shear viscosity, *ζ* the active contraction exerted by the contractile myosin motors. *ρ*_*m*_ and *ρ*_*a*_ are the density of the myosin motors and the actin network specified below.

#### Mechanics of the IF network

To model the mechanics of the IF network and obtain its velocity field *v*^*IF*^, we define an analogous equation for balance of linear momentum^112^ along with boundary conditions

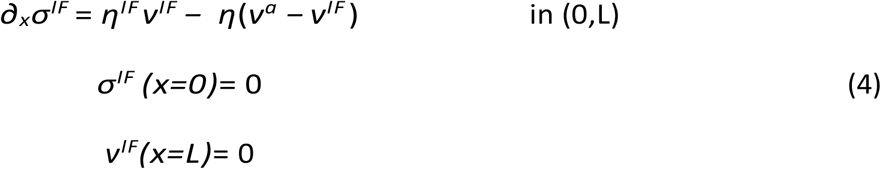

where we impose zero velocity at the nucleus^53^, which is also consistent with results (main text Fig. 4l). We impose that the network is traction-free at the cell front.

As for the actin network, the first term in the right-hand side represents the friction between the IF network and the ECM, which we assume to be proportional to the IF network velocity with friction constant *η*^*IF*^. Following the same argument as for the interaction of the F-actin network with the ECM, we assume the same space dependency of *η*^*IF*^, consistent with the adhesion distribution observed in the in-vitro system (Method Fig. 1c). Note, however, that this spatial dependence had a minor effect on the results. The second term is the friction resulting from inter-network relative motion.

We model the constitutive response of the IF network as a viscoelastic solid obeying:

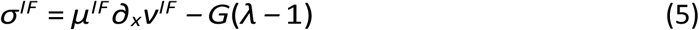

where the first term accounts for the viscosity of the IF network with shear viscosity *μ*^*IF*^ and the second term is an elastic stress, where *G* is the bulk elastic modulus and λ is the stretch ratio of the network. Based on previous data, we consider *G* to be proportional to the density of the IF network as *G* = *G*_0_*ρ*^*IF*^ ^49^, where *G*_0_ is the elastic modulus at unit normalized density. The elastic modulus of the keratin network is also assumed to increase as the crosslinking of keratin network increases^62,115^. Through hemidesmosomes, keratin network attachment to the substrate will increase network crosslinking^60^. To introduce this aspect in the simplest possible way, we vary *G*_0_ proportionally to substrate friction, which also depends on hemidesmosomes, following 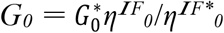. We choose 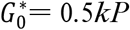 at a reference friction *η^IF*^_0_ = 5* < 0 kPa·s/μm^2^ such that the value of *G*_0_ in the WT and *β*4R1281W cases fits the respective bulk elastic moduli of the experimental data (main text Fig. 4p,q). Of note, experimental bulk moduli capture contributions of all cytoskeletal networks and not only IFs. In our model, we assign these values to the IF network for simplicity due to its major role. Importantly, the differences between the WT and mutant conditions (which are the main focus of the model) are indeed caused specifically by changes in the IF network.

The relation between network stretch and velocity is given by the kinematic relation ∂_t_λ + ∂_x_(v^IF^λ) = 0 ^112^. However, to model the fact that elastic stresses dissipate over time, e.g. as a result of turnover, or equivalently that the elastic strain λ-tends to zero as the network is rebuilt with a rate constant *τ*^*λ*^, we consider the following previously considered advection-reaction equation for the evolution of λ ^112^

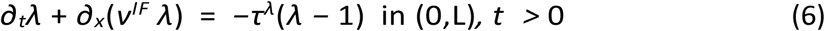

#### Model of F-actin, myosin and IF density

To describe the distribution in time- and space of the intracellular F-actin, myosin and IF densities, we consider three transport advection-diffusion reaction equations. We model the effective transport of the actin network density, *ρ*^*a*^, as:

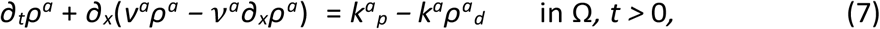

where *v*^*a*^ is the diffusion constant, and *k*^*a*^_*p*_ and *k*^*a*^_*d*_ are the polymerization and depolymerization rates of the network, respectively. We impose *k*^*a*^_*p*_ = *k*^*a*^_*d*_ to normalize the density to one. We impose homogeneous natural boundary conditions at both sides of the domain.

Similarly, we model the effective transport of the IF network density, *ρ*^*IF*^, as:

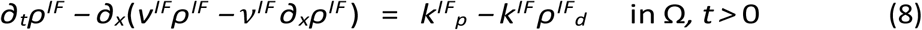

where *v*^*IF*^ is the diffusion constant, and *k*^*IF*^_*p*_ and *k*^*IF*^_*d*_ are the polymerization and and depolymerization rates of the IF network. We impose *k*^*IF*^_*p*_ = *k*^*IF*^_*d*_ to normalize the density to one at steady state. We impose again zero flux boundary conditions at each cell end to describe that the IF network cannot enter or leave the cell membrane in the front of the cell and to reflect symmetry boundary conditions at the nuclear side.

Finally, we model the effective transport of myosin motor density bound to the F-actin network, *ρ^m^*, as a transient advection-diffusion problem^116^:

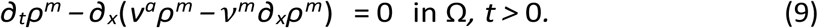

This equation reflects the assumption of a much faster attachment rate of motors to the F-actin network than the detached rate into the unbound cytosolic form. Effectively, bound myosin motors are advected with velocity *v^a^* ^57,116,117^ and diffuse with diffusion constant *vm*. Again, we impose zero flux boundary conditions at both ends.

#### Model of nuclear deformation and sphericity

The composite network model described above allows us to compute the total stress at the nucleus by evaluating *σ^a^* + *σ^IF^* at x = 0. We obtain a total stress of 0.1kPa and 0.11kPa for the control and mutant cases, respectively. We model the nucleus as a linear element so that nuclear strain ε is given by *σ^IF^* + *σ^a^* =G^N^ε, where G^N^ is the nuclear elastic modulus taken from Fig. 4Q. Given ε, and hence the in-plane linear stretch ε = 1+ε, and assuming equibiaxial strain and nuclear incompressibility, we compute the out-of-plane linear stretch as 1/ε2 = 1/(1+ε)^2^. Thus, an initially spherical nucleus of radius R ends up as an oblate ellipsoid with in-plane semiaxes R(1+ε) and out-of-plane semiaxis R/(1+ε)^2^, which provides an estimate for nuclear shape and allows us to compute sphericity. We obtain a sphericity of 0,98 and 0,81 for control and mutant cells.

#### Modelling intracellular network dynamics in WT cells

To verify our model, we first simulate the dynamics of the intracellular networks in the WT cells and validate the results against the experimental data. Our computational model includes a number of model parameters that do not change as we change conditions of this study. These model parameters are presented in Table 1. Note that many of these model parameters are obtained from the literature, although we allowed for slight variations to better fit experimental results. Regarding the diffusion parameters of the actin and keratin networks, we assumed that they do not diffuse and are transported mainly because of convection. Therefore, we introduced small diffusive terms to stabilize the numerical solution of the equations. For the turnover rates of the system, we assumed that the attachment of free myosin is fast, so that the effective transport equation for the attached myosin motors is given as in Eq. 9. We assumed that actin turnover is 2 orders of magnitude faster than that of the keratin network.

**Table 1:** Material parameters of the computational model of the acto-myosin and IF network. All references were used to obtained magnitudes of the corresponding parameters. * indicates that the values were fitted in this work.

|  | Actin |  | IF |  |
| --- | --- | --- | --- | --- |
|  | Value | Ref. | Value | Ref. |
| $\mu$ [kPa·s] | 30 | 116,118 | 10 | 112 |
| $\eta$ [kPa·s/μm] | 8 | 116,118,119 | 8 | * |
| $\zeta$ [kPa] | 0.1 | 57,118 | - | - |
| $G_0^*$ [kPa] | - | - | 0.5 | * |
| $k_p / k_d$ | 0.1/0.1 | 120,121 | 0.001/0.001 | * |
| $v$ [μm/s] | 0.001 | * | 0.001 | * |
| $\eta$ [kPa·s/μm] | 4 | * | 4 | * |
| $a$ [-] | 4 | * | 4 | * |

Our results for WT cells show an accumulation of IFs close to the nucleus while F-actin density is more uniformly distributed along the cell, both in agreement with experimental results (main text Fig. 4f,g). Actin flow is powered by the active pulling of myosin motors that has to balance the friction with the ECM and with the IFs (Methods Fig. 1b). The actin retrograde flow decays quickly from 1.5 nm/s at the cell front to zero at the nucleus (main text Fig. 4h). The IF network flow does not have intrinsic activity in our model and is dragged by the motion of the actin network and experiences the friction with the ECM. The velocity of IFs follows a similar decay from the cell periphery to the nucleus, but it is 1 order of magnitude slower than actin flow (main text Fig. 4i). As a result of these dynamics, the IF network accumulates close to the nucleus and imposes a viscous stress to the cell nucleus.

Results also show that the actin network progressively goes from a compressive state of −0.27 kPa, close to the cell membrane, to a tensional state of 0.11 kPa at the nucleus. The IF network is elastically in compression in the entire cell domain, except for a small region close to the cell membrane where it is stretched. At the nuclear region, IFs exert a compressive stress of 0.01kPa. Therefore, total tension applied to the cell nucleus is 0.1kPa. The frictional forces on the ECM, resulting from actin and IF flows, are high at the cell periphery, as observed experimentally. It is also important to note that the frictional forces applied by the actin network through FAs, are higher (1 order of magnitude) than the frictional forces that the IF network applies on the ECM. Again, this is in close agreement with the experimental observations (no change in traction forces, main text Supplementary Fig. 3e,f).

#### Modelling intracellular network dynamics in β4R1281W mutant cells

We then simulate the network dynamics in *β*4R1281W mutant cells. We assume that the main change in the system is a reduction in the friction of IFs with the substrate, and therefore we lower *η^IF^* from 8 to 2 kPa s/μm^2^. Mutant cells also have decreased keratin-substrate cross-linking and thereby decreased elasticity, which as explained above is modelled by introducing a linear relation between *η*^*IF*^_0_ and *G*_0_.

Our results show that the mutant condition leads to negligible effects in the density and velocity of the actomyosin cytoskeleton, and in the corresponding traction forces on the substrate. There is a small increase in nuclear stress (8%), that occurs because reduced IF-substrate friction decreases the stiffness of the IF network and, therefore, reduces the compressive forces applied onto the nucleus. However, the main effects in nuclear strain are not explained by this small increase in stress, but from a decrease in the elasticity (stiffness) of the IF network around the cell nucleus. IFs are crosslinked to the nucleus and contribute to its mechanical properties^48,62,115^, and thus according to the model the overall stiffness of the nucleus and its surrounding cytoskeleton should decrease, as verified experimentally (main text Fig. 4n,p,q). Overall, the combined effects of increased stress and decreased stiffness lead to an increase in nuclear strain for the mutant cells, promoting nuclear mechanotransduction.

## Notes

### Competing Interest Statement

The authors have declared no competing interest.

