## Supplementary Material for "The laminin-keratin link shields the nucleus from mechanical deformation and signalling"

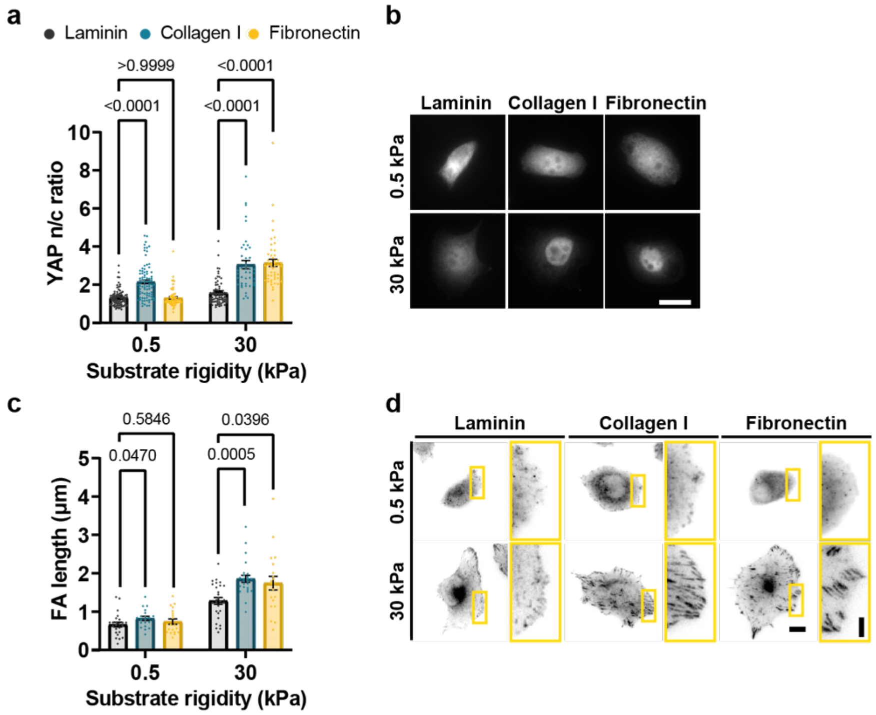


**Supplementary Fig. 1| Rigidity response of myoepithelial cells on different substrates.**

**a.** Quantification of n/c YAP ratio ($n$ = 82/84/61, 70/43/48 cells for laminin/collagen/fibronectin for 0.5 and 30 kPa respectively; mean of at least 2 independent experiments). The effect of both rigidity and substrate coating is significant. ($P$ < 0.0001, two-way ANOVA, Bonferroni correction for multiple comparison).

**b.** Sample YAP images for human myoepithelial cells on laminin/collagen/fibronectin substrates of 0.5 and 30kPa stiffness; scale bar is 20 µm.

**c.** Quantification of focal adhesion (FA) length from phospho-paxillin imaging ($n$ = 24/20/20, 28/25/20 cells for laminin/collagen/fibronectin for 0.5 and 30 kPa respectively; mean of at least 2 independent experiments). The effect of both rigidity ($P$ < 0.0001) and substrate coating is significant. ($P$ =0.0002, two-way ANOVA, Bonferroni correction for multiple comparison).

**d.** Sample phospho-paxillin images for human myoepithelial cells on laminin/collagen/fibronectin substrates of 0.5 and 30kPa stiffness; scale bars are 10 µm (main images)/ 4 µm (zoomed images).

Error bars represent mean ± s.e.m.

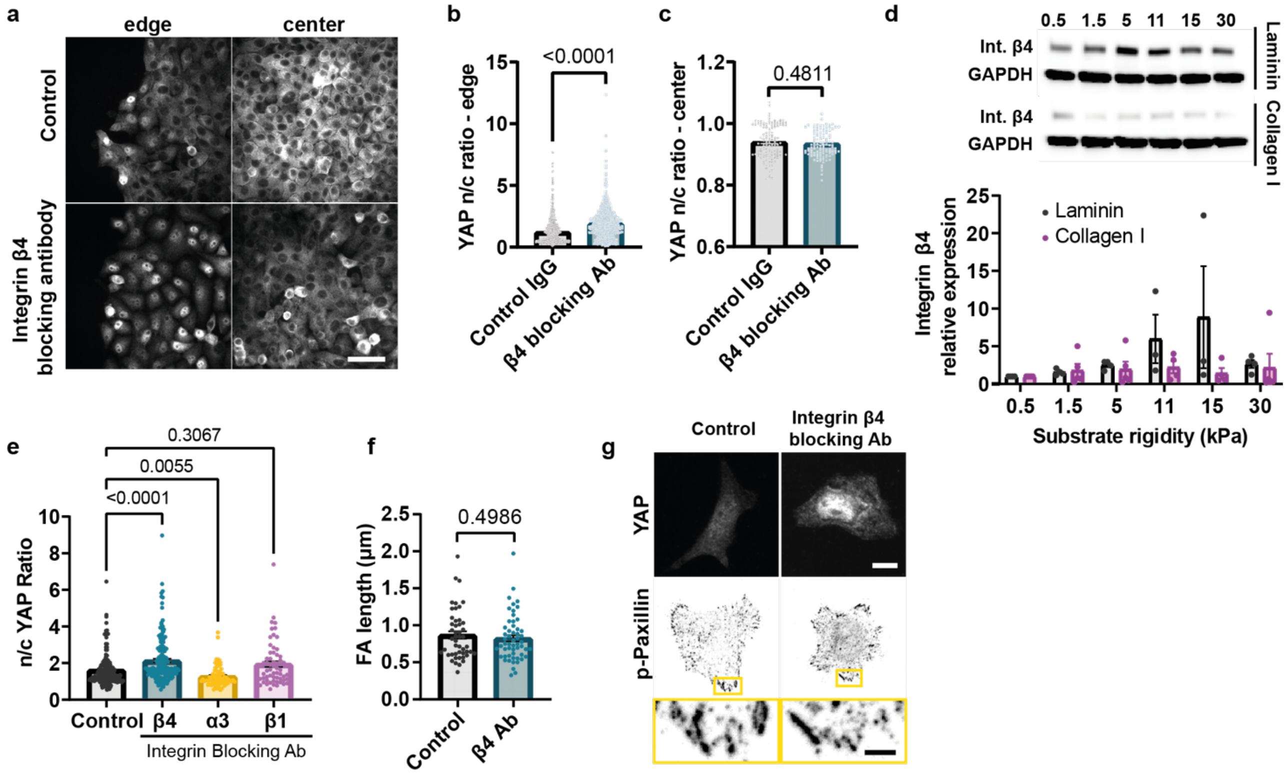


**Supplementary Fig. 2| Further experiments on the role of α6β4 integrins.**

- **a.** YAP stainings of MCF10A cells treated with control IgG/β4 blocking antibodies for cells grown on 11kPa substrate stiffness. Left panel shows cells at the edge of the monolayer and on the right corresponding images of the monolayer centre; scale bar 50µm.
- **b.** Quantification of n/c YAP ratios of MCF10A cells at the edge of the monolayer upon treatment with control IgG or integrin β4 blocking antibody ($P$<0.0001, two-tailed Mann-Whitney test, n= 841/723 cells from 3 independent experiments).
- **c.** Quantification of n/c YAP ratios of MCF10A cells at the centre of the monolayer upon treatment with control IgG or integrin β4 blocking antibody ($P$>0.05, two-tailed Mann-Whitney test, n=128/117 cells from 3 independent experiments).
- **d.** Western blot for integrin β4 expression levels of MCF10A cells on laminin or collagen coated gels of different rigidities, quantification of at least 3 independent experiments per condition.
- **e.** Average values of nuclear/cytosolic YAP ratio of myoepithelial cells seeded on laminin coated PAA gels of 30kPa stiffness upon treatment with control or integrin β4/α3/β1 blocking antibodies ($n$ = 145/165/76/61 cells for control/ β4/α3/β1; mean of at least 3 independent experiments). The effect of integrin β4 and α3 blocking is significant. ($P$ < 0.0001, Kruskal-Wallis test, Dunn's correction for multiple comparison).
- **f.** Focal adhesion length from pPAX stainings of control and integrin β4 blocking Ab treated myoepithelial cells (n= 48/55; mean of 3 independent experiments). No significant effect was observed ($P$>0.05, two-tailed unpaired t-test).

**g.** Corresponding images of YAP and phospho-Paxillin stainings for control and integrin β4 blocking Ab treated myoepithelial cells; scale bars are 10 µm (main images)/ 2.5 µm (zoomed images). Error bars represent mean ± s.e.m.

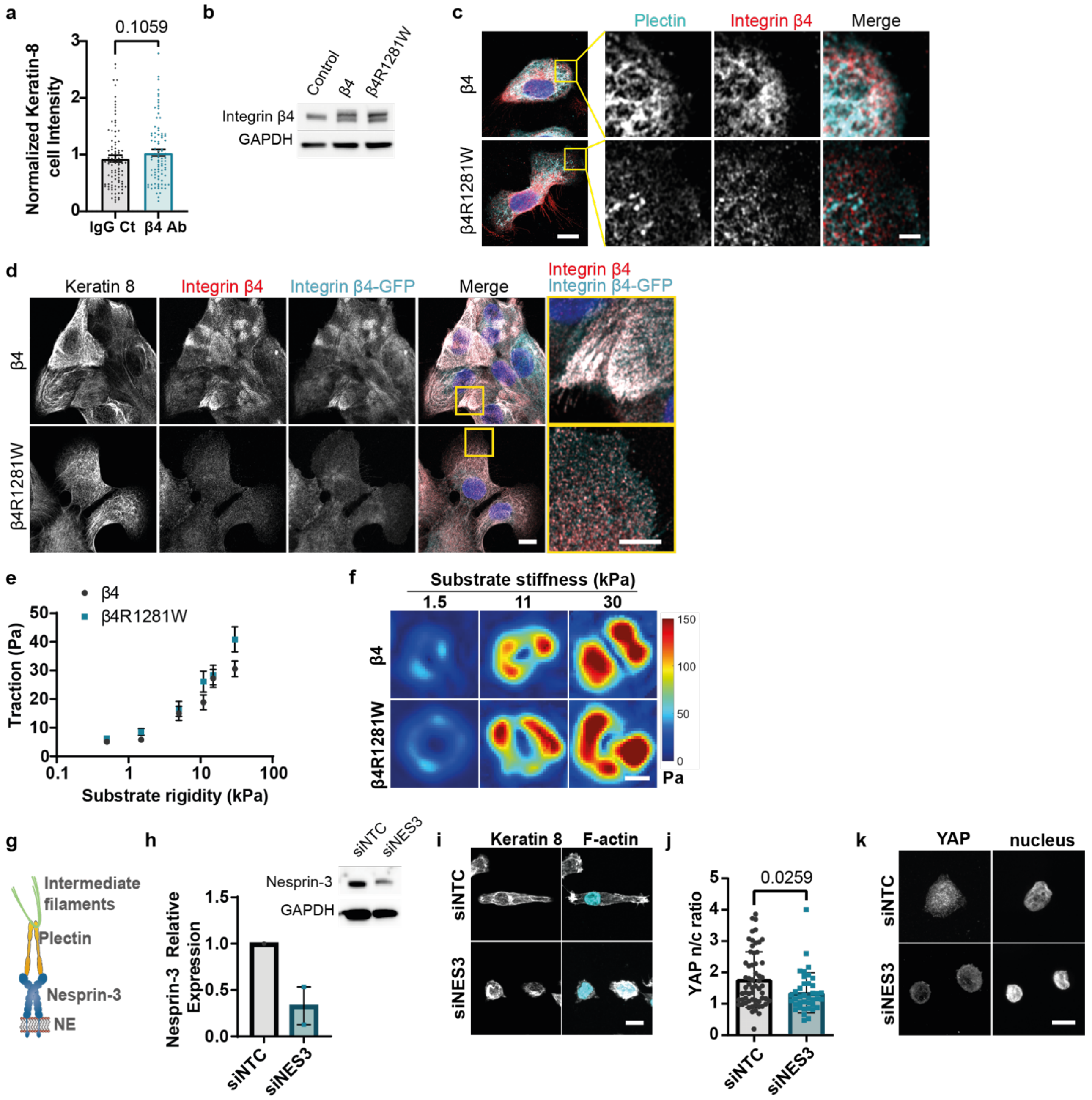


**Supplementary Fig. 3| Further characterization of the effects of keratin-laminin links.**

- **a.** Mean intensity of sum confocal projections of keratin-8 stainings ($p>0.05$, unpaired two-tailed t-test, n= 93/89 cells for control (IgG Ct) or integrin β4 blocking conditions (β4 Ab) for 6 independent experiments). Error bars represent mean ± s.e.m.
- **b.** Western blot of integrin β4 in MCF10A cells in control conditions or after overexpressing WT or β4R1281W β4 in MCF10A cells.
- **c.** Stainings for plectin (cyan) and integrin β4 (red) in cells overexpressing WT integrin β4 or β4R1281W integrin β4; scale bars 10 µm (main images) and 2 µm (zoomed images).

**d.** Stainings for keratin 8, integrin β4 and GFP (overexpressed integrin) for cells overexpressing WT integrin β4 or β4R1281W; scale bar is 10 µm. Zoomed insert shows total integrin β4 (red) and GFP (cyan) stainings; scale bar is 5µm.

**e.** Tractions for WT integrin β4 or β4R1281W overexpressing cells. (*n* = 30/41, 30/29, 30/24, 31/28, 34/29, 40/32 cells for β4/β4R1281W and increasing rigidity; mean of 3 independent experiments). The effect of rigidity is significant. ($P < 0.0001$, two-way ANOVA). Error bars represent mean ± s.e.m.

**f.** Corresponding traction maps of WT or R1281W integrin β4 overexpressing cells; scale bar is 10µm.

**g.** Schematic representation of the connection of nesprin-3 to intermediate filaments through plectin; NE=nuclear envelope.

**h.** Western blot for cells transfected with a control non-targeting siRNA (siNTC) or nesprin-3 siRNA (siNES3) and quantification (n=2 independent repeats). Error bars represent mean ± S.D.

**i.** Keratin-8 and Phalloidin (F-actin) stainings for control and Nesprin-3 KD MCF10A cells; scale bar is 10 µm.

**j.** Quantification of YAP n/c ratios for non-targeting (siNTC) or nesprin-3 siRNA (siNES3) transfected MCF10A cells ($P$=0.0259, unpaired two-tailed t-test, *n* = 60/39 control/siRNA transfected cells respectively, from 2 independent experiments). Error bars represent mean ± s.e.m.

**k.** Sample YAP stainings for siNTC or siNES3 transfected cells; scale bar is 10 µm.

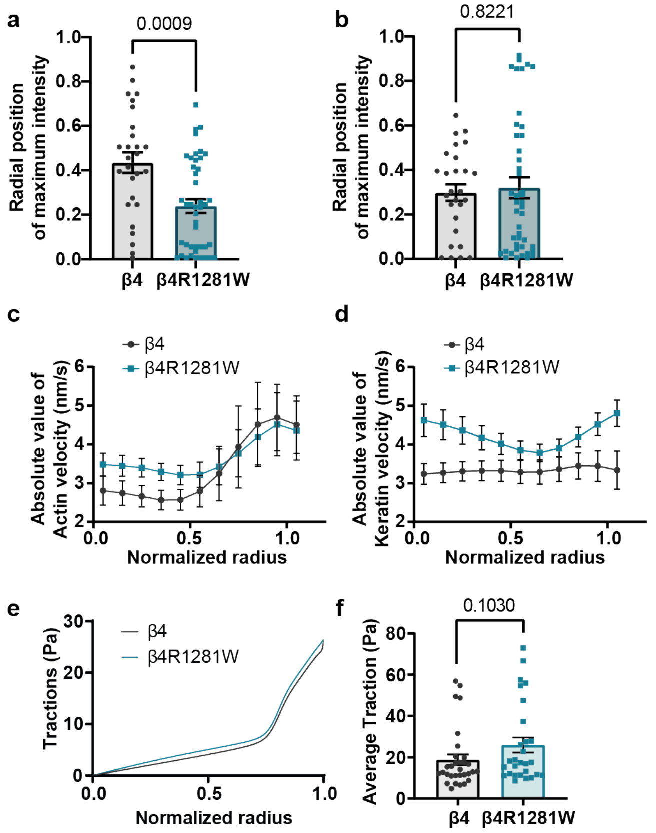


**Supplementary Fig. 4| Further characterization of differences between integrin β4 or β4R1281W overexpressing cells and model predictions.**

**a.** Plots showing the radial position of maximum keratin-8 signal for β4 or β4R1281W overexpressing cells. R=1, cell periphery, R=0, cell centre (*P*=0.0009, Mann-Whitney test, *n* = 27/43 cells for β4 and β4R1281W).

**b.** Plots showing the radial position of maximum actin signal. (*P*>0.05, Mann-Whitney test, *n* = 27/43 cells for β4 and β4R1281W).

**c.** Experimental quantifications of absolute actin retrograde flows along the cell radius (cell periphery R=1 and centre R=0). The combined effect of β4 mutation and radial actin velocities is not significant (*P* = 0.6927, two-way repeated measures ANOVA, *n* = 12/12 cells for β4 and β4R1281W).

**d.** Experimental quantifications of absolute keratin-18 retrograde flows along the cell radius. The combined effect of β4 mutation and radial keratin velocities is significant (n=12, *P* = 0.0073, two-way repeated measures ANOVA, *n* = 12/12 cells for β4 and β4R1281W).

**e.** Model prediction for radial tractions.

**f.** Experimental quantification of average cell tractions at 11kPa from Supplementary Fig. 3e (*P* > 0.05, two-tailed Mann Whitney test, n=31/28 cells for β4/β4R1281W).

Error bars represent mean ± s.e.m.

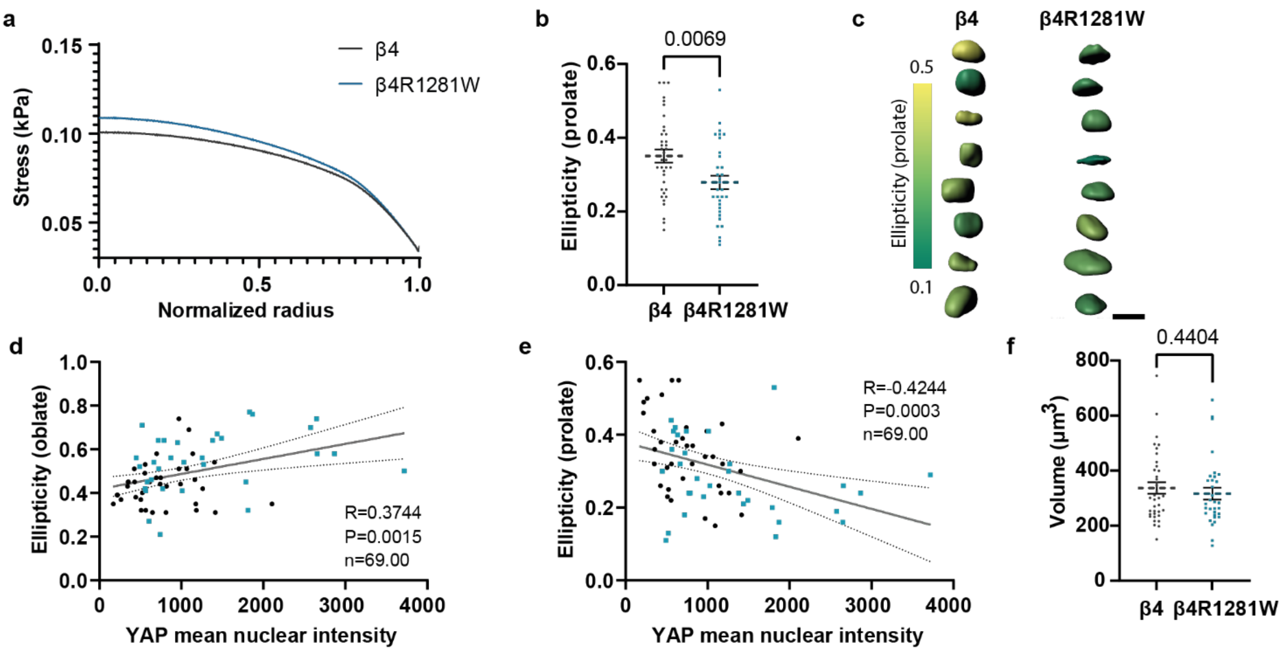


**Supplementary Fig. 5| Further characterization of nuclear shape.**

- **a.** Model prediction for radial cell stress. Stress applied to the nucleus is the point corresponding to radius = 0.
- **b.** Nuclear ellipticity (prolate) measurements (*P*=0.0069, two-tailed unpaired t-test, n=37/32 for β4/β4R1281W in 3 independent experiments).
- **c.** 3D segmentation and ellipticity (prolate) colour-coded nuclei for integrin β4 (left) and β4R1281W (right) overexpressing cells; scale bar is 10 µm.
- **d.** Correlation between nuclear YAP intensity and ellipticity for both β4 (black circles) and β4R1281W (blue squares) integrin expressing cells (R is Spearman's correlation coefficient).
- **e.** Correlation between Nuclear YAP intensity and ellipticity (prolate) for both β4 (black circles) and β4R1281W (blue squares) integrin expressing cells (R is Spearman's correlation coefficient).
- **f.** Nuclear volume measurements (*P*>0.05, two-tailed unpaired t-test, n=37/32 for β4/β4R1281W in 3 independent experiments).

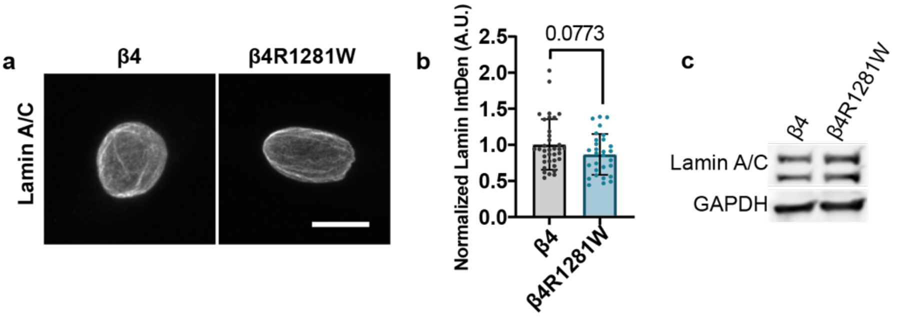


**Supplementary Fig. 6| Lamin A/C quantification in integrin β4 or β4R1281W expressing cells.**

**a.** Lamin A/C stainings; scale bar is 10 µm.

**b.** Quantification of normalized Integrated Density (*P*>0.05 two-tailed unpaired t-test n=35/29 for β4/β4R1281W from 3 independent experiments).

Western blot for Lamin A/C for integrin β4 and β4R1281W overexpressing cells (n=2).

**Supplemental items information**

**Supplementary Video 1| Actin and keratin velocities for wt integrin β4 expressing cells.**

Spinning disc confocal time-lapse imaging of a cell co-transfected with Lifeact-GFP and Keratin-18-mCherry, seeded on a laminin-coated circular pattern. Scale bar is 20µm.

**Supplementary Video 2| Actin and keratin velocities for integrin β4R1281W expressing cells.**

Spinning disc confocal time-lapse imaging of a cell co-transfected with Lifeact-GFP and Keratin-18-mCherry, seeded on a laminin-coated circular pattern. Scale bar is 20µm.
